# *Drosophila* Tropomodulin is required for multiple actin-dependent processes in myofiber assembly and maintenance

**DOI:** 10.1101/2022.08.05.502981

**Authors:** Carolina Zapater i Morales, Peter J. Carman, David B. Soffar, Stefanie E. Windner, Roberto Dominguez, Mary K. Baylies

**Affiliations:** Biochemistry, Cell & Developmental Biology, and Molecular Biology (BCMB) program, Weill Cornell Graduate School of Medical Sciences, New York, NY 10065; Developmental Biology Program, Sloan Kettering Institute, Memorial Sloan Kettering Cancer Center, New York, NY 10065; Biochemistry and Molecular Biophysics Graduate Group, Perelman School of Medicine, University of Pennsylvania, Philadelphia, PA 19104; Department of Physiology, Perelman School of Medicine, University of Pennsylvania, Philadelphia, PA 19104

**Keywords:** Skeletal muscle, Tropomodulin (Tmod), *Drosophila*, sarcomere length, myofibril orientation, tension-mediating proteins, misshapen nuclei

## Abstract

Proper muscle contraction requires the assembly and maintenance of sarcomeres and myofibrils. While the protein components of myofibrils are generally known, less is known about the mechanisms by which they individually function and together synergize for myofibril assembly and maintenance. For example, it is unclear how the disruption of actin filament (F-actin) regulatory proteins leads to the muscle weakness observed in myopathies. Here, we show that knockdown of *Drosophila* Tropomodulin (Tmod) results in several myopathy-related phenotypes, including reduction of muscle cell (myofiber) size, increased sarcomere length, disorganization and misorientation of myofibrils, ectopic F-actin accumulation, loss of tension-mediating proteins at the myotendinous junction, and misshaped and internalized nuclei. Our findings support and extend the tension-driven self-organization myofibrillogenesis model. We show that, like its mammalian counterpart, *Drosophila* Tmod caps F-actin pointed-ends, and this activity is critical for cellular processes in different locations within the myofiber that directly and indirectly contribute to the maintenance of muscle function. Our findings provide significant insights to the role of Tmod in muscle development, maintenance, and disease.

**SUMMARY STATEMENT:** *Drosophila* Tropomodulin knockdown in larval myofibers results in myopathy-related phenotypes. Our findings support that Tmod acts in actin-related processes at different subcellular locales, all critical for muscle integrity and function.

## INTRODUCTION

Skeletal muscle cells (myofibers) are essential for survival. Myofibers are comprised of myofibrils, each consisting of arrays of repeating units known as sarcomeres, the muscle’s fundamental contractile machinery. These contain three main structural elements: thin filaments, thick filaments, and Z-discs [reviewed in Sweeney and Hammers (2018)]. Z-discs are the thin filament attachment site and are composed of many proteins (e.g.: α-actinin and Zasp). Thin filaments are comprised primarily of actin filaments (F-actin) and actin binding proteins, including proteins important for anchorage to the Z-disc (e.g.: CapZ), for contraction (e.g.: troponin and tropomyosin), and for F-actin length regulation (e.g.: nebulin and Tmod) [reviewed in Szikora et al. (2022), Ghosh and Fowler (2021), Fowler and Dominguez (2017)]. Thick filaments are composed primarily of myosin and myosin-associated proteins. Thin filaments sliding past thick filaments brings Z-discs closer together (contraction), generating force in a calcium-regulated manner. While much is known about protein composition and function of sarcomeres and myofibrils, the mechanisms controlling their formation and maintenance remain elusive.

Although myofibrillogenesis is not well understood, the most widely accepted model suggests tension is essential. According to this model, once the myofiber has attached to tendon cells, short F-actin stochastically attach to each other and to the myofiber ends via tension-activated proteins such as β-Integrin and Talin. Through a positive feedback loop, initial tension attracts more tension-activated proteins, leading to further tension and myofibril assembly and maturation along the myofiber’s longitudinal axis (Lemke and Schnorrer, 2017, Lemke et al., 2019, Pines et al., 2012, Weitkunat et al., 2014, Sparrow and Schock, 2009). Subsequently, additional growth and organization along myofibrils also require tension (LeGoff and Lecuit, 2015). Several other processes regulate myofibrillogenesis and myofibril maintenance. Firstly, mitochondria influence myofibrillogenesis and vice versa via the mechanical interplay between mitochondrial dynamics and myofibril morphogenesis (Avellaneda et al., 2021). Secondly, myofibrils undergo protein turnover without compromising their organization during contraction and myofibrillogenesis (Sanger and Sanger, 2008, Ono, 2010, Littlefield and Fowler, 2008). Finally, actin regulatory proteins that control monomeric (G-actin) to F-actin ratios in the cell are critical for myofibril assembly and turnover (Littlefield and Fowler, 2008, Deng et al., 2021). While contributing factors to myofibril formation and maintenance have been identified, their interplay and molecular mechanisms are incompletely understood.

Mutations in sarcomere genes are linked to skeletal muscle diseases such as nemaline myopathy (NM), which presents with muscle weakness (de Winter and Ottenheijm, 2017, Sewry et al., 2019). NM is associated specifically with mutations in thin filament genes, including ACTA1, NEB, LMOD3, CFL2, TPM2, TPM3, TNNT1, TNNT3 and MYPN [reviewed in Laitila and Wallgren-Pettersson (2021), Christophers et al. (2022)]. In addition to the growing list of mutations identified in human patients, Tmod mutant zebrafish show a NM-like phenotype (Berger et al., 2014). Studies in model systems have demonstrated that Tmod regulates F-actin length by capping pointed-ends to inhibit monomer addition/dissociation (Molnar et al., 2014, Berger et al., 2022, Littlefield et al., 2001). Tmod is related in sequence and structure to Leiomodin (Lmod), an F-actin nucleator [(Chereau et al., 2008) and reviewed in Fowler and Dominguez (2017), Yamashiro et al. (2012)], whose deficiency has been linked to NM (Yuen et al., 2014, Cenik et al., 2015). Some Tmods also have low nucleation activity (Fischer et al., 2006, Yamashiro et al., 2010, Yamashiro et al., 2014). While humans have four *TMOD* and three *LMOD* genes that cap and nucleate F-actin, respectively, *Drosophila* has no Lmod, but expresses 18 Tmod splice variants. It is unknown, however, which Tmod isoforms are most abundant in *Drosophila* muscle and whether individual isoforms cap or nucleate F-actin. It is also unknown whether and how the disruption of Tmod isoforms in *Drosophila* would result in the NM-like defects observed in other models.

Previous muscle research focused on the role of Tmod and Lmod in regulating sarcomeric F-actin. However, similar to other cell types, cytoplasmic F-actin plays several essential roles in myofibers: maintenance of intracellular structure by tethering myofibrils to the plasma membrane and extracellular matrix (ECM), organization of myofiber organelles and vesicle trafficking, and movement and anchoring of myonuclei at the myofiber periphery [reviewed in Davidson and Cadot (2021), Charvet et al. (2012), Kee et al. (2009)]. The disruption of cytoplasmic F-actin dynamics could contribute to muscle deterioration and weakness and has not been comprehensively studied. Here, we analyze the effects of Tmod knockdown (Tmod-KD) in *Drosophila* larval muscle. We observe progressive muscle deterioration structurally and functionally, with defects in muscle size, sarcomere length, myofibril homeostasis, integrin-based myofibril attachment, and myonuclear shape and activity. Alongside with myofibrillar organization breakdown, Tmod-KD myofibers show ectopic F-actin accumulations. Transcriptomic analyses reveal downregulation of proteasomal degradation and upregulation of oxidative metabolism. Biochemically, *Drosophila* Tmod exclusively acts as an actin capping protein which, in addition to regulating thin filament length, is involved in establishing tension along the myofiber. The phenotypes generated by Tmod loss support and extend the leading hypothesis of myofibrillogenesis and suggest a role for Tmod in myofibril assembly and maintenance. We propose that Tmod’s function as an F-actin capping protein is critical in a variety of locations within the myofiber that directly and indirectly maintain muscle function.

## RESULTS

### *Drosophila* Tmod is required for larval muscle integrity

To identify genes with a critical role in muscle development and disease, we performed a limited screen in *Drosophila melanogaster*. We used a muscle-specific driver (*Dmef2-Gal4*) and available RNAi lines to knock down homologues of human proteins implicated in myofibril/sarcomere structure and function (Schnorrer et al., 2010) (Table S1). We screened for embryonic lethality, muscle defects in late third instar larvae, pupal lethality, and adult flight (Table S2). Expression of *tmod*RNAi resulted in phenotypes that suggested Tmod maintains muscle integrity and function. We confirmed the KD phenotypes using three different RNAi lines and a homozygous *tmod* mutant, and assessed KD levels by qRT-PCR, immunofluorescence, and Western blot (Fig. S1). Based on these results, we selected *tmod* for further study and used one RNAi line, dsRNA-HMS02283 (Perkins et al., 2015), for subsequent KD experiments as it exhibited an intermediate phenotype and degree of mRNA KD.

*Drosophila* larvae are segmented organisms with three thoracic and eight abdominal body segments (A1-A8), each consisting of ∼30 unique body muscles (Bate, 1990). Here, we focused on two ventral longitudinal (VL) muscles, VL3 and VL4 (also named muscles 6 and 7), which are well characterized, easily assessable, flat rectangular cells with nuclei located on one cell surface (Fig. 1A-C) (Windner et al. (2019). Muscle-specific Tmod-KD resulted in a striking phenotype at the end of larval development, with a significant reduction in myofiber size (Fig. 1D,E). While the myofiber length was conserved, the width was reduced by 30-60% (Fig. 1D-F). Furthermore, we observed several distinct morphological features: increased sarcomere length (Fig. 1G), disorganized, misoriented myofibrils (Fig. 1I), ectopic F-actin accumulation (Fig. 1D,E), and misshaped, internalized nuclei (Fig. 1J). Strikingly, intact myofibrils with changes in sarcomere length were restricted to the myofiber’s cuticular side. In contrast, the myofiber’s visceral surface, which includes the myonuclei, was disorganized (Fig. 1E). These observations supported a crucial function for Tmod in regulating sarcomeric actin and myofibril stability. Furthermore, they suggested additional Tmod role(s) in the regulation of cytoplasmic actin, including nuclear stabilization and other cellular processes.

**Fig. 1:**
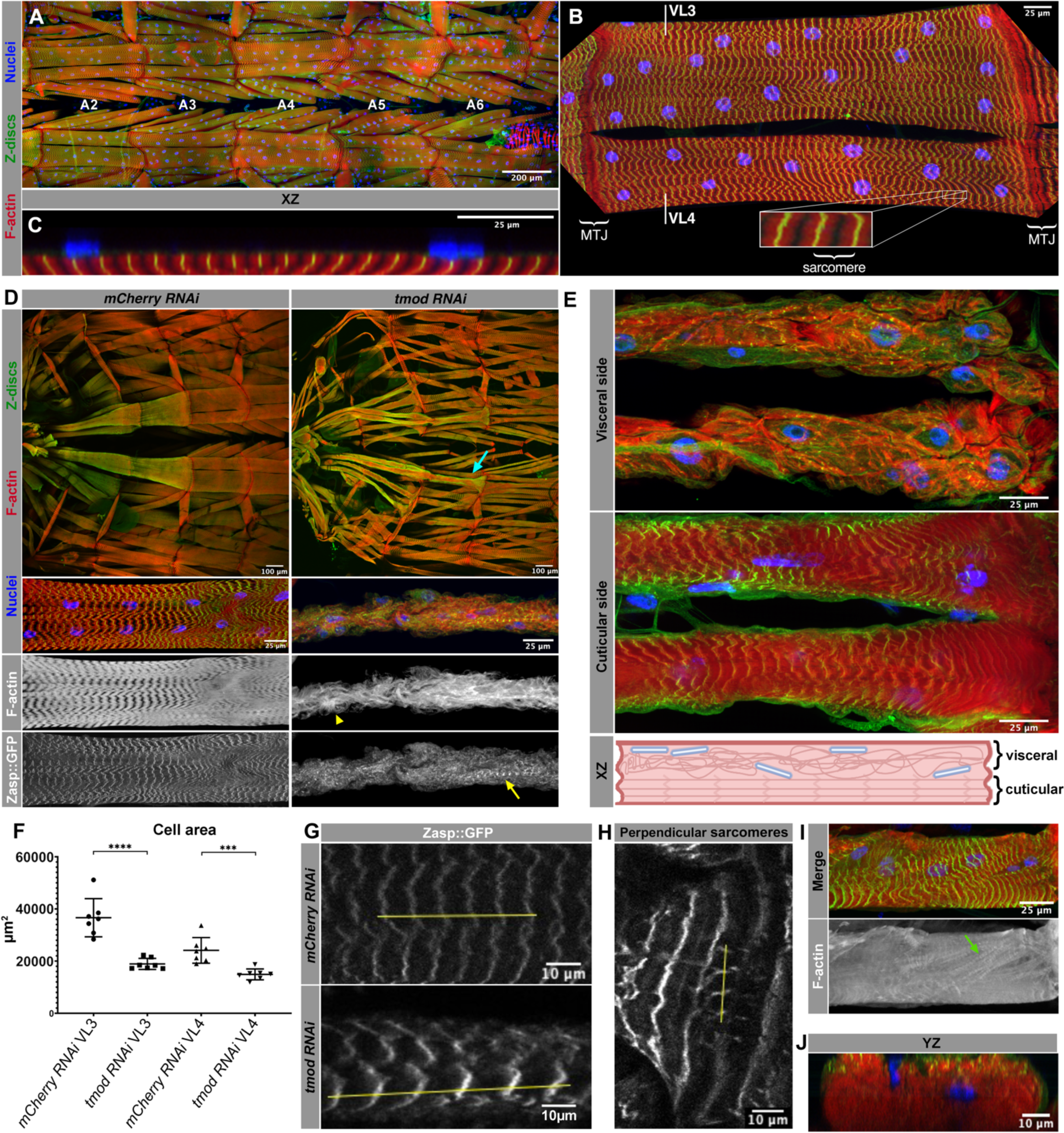
Tmod is required for larval muscle integrity. (A) Third instar *Drosophila* larval filet, abdominal segments (A2-6) and body wall muscles (red, phalloidin) with Z-discs (green, Zasp::GFP) and nuclei (blue, Hoechst); anterior, left. (B) VL3 and VL4 muscles showing MTJs and zoomed-in panel of a sarcomere (Z-stack projection). (C) VL4 myofiber longitudinal cross-section with nuclei on the cell surface. (D) Overview of anterior musculature in control (*mCherryRNAi*) and Tmod-KD (*tmodRNAi*) larvae (top). Higher magnification of VL3 muscle merge, F-actin and Zasp::GFP (bottom 3 panels). Cyan arrow indicates width reduction of the myofiber. Yellow arrowhead: ectopic F-actin. Yellow arrow: remaining myofibril. (E) Visceral view of VL3 and VL4 Tmod*-*KD myofibers showing disorganized myofibrils and ectopic F-actin (top), cuticular view showing conserved sarcomeres (middle) and cross-sectional diagram showing integration of both visceral and cuticular sides in a myofiber (bottom). (F) VL3 and VL4 myofiber areas (A2) in control and Tmod-KD wandering late third instars. Graph represents one experiment (N>3 replicates, each with n_VL3_=7 myofibers and n_VL4_=7 myofibers per genotype; p_VL3_<0.0001 and p_VL4_=0.0006, two-tailed unpaired Student’s t-test). Mean ± SD. (G) Sarcomere length in control and Tmod-KD myofibers. Yellow line spans six Z-discs. (H) Perpendicular sarcomeres in Tmod-KD myofibers. Yellow line spans four Z-discs. (I) Disorganized, misoriented myofibrils. Green arrow indicates diagonal myofibrils. (J) Optical cross-section of Tmod-KD myofiber displaying internalized and misshapen nuclei. Scale bars: 200μm (A), 100μm (top panels in D), 25μm (B, C, bottom 3 panels in D, E, and I) and 10μm (G, H, and J).

### Tmod-KD myofibers deteriorate in a specific pattern

*Drosophila* larval development can be divided into four instar stages during which body wall muscles grow dramatically in size (25 to 40-fold): first, second, early third, and late third instars (the last two hours of which is called the “wandering” stage). Over this four-day period, new sarcomeres and myofibrils are added, while nuclear number remains constant (Demontis and Perrimon, 2009, Bai et al., 2007, Balakrishnan et al., 2020). While muscle phenotypes were found in all Tmod-KD late third instars (see also Fig. S1B), not all segments were affected equally. Instead, we found a gradient of muscle phenotypes along the larval anterior-posterior (AP) axis. Myofibers in more anterior segments (A1, A2) had severely disrupted myofibrils and nuclei, while posterior ones (A5, A6) maintained nuclear shape and position and exhibited a milder myofibrillar phenotype.

We analyzed Tmod-KD phenotypes along the larval AP axis throughout larval development. In first instars, we observed wildtype (WT)-like muscles, which contained the typical conserved arrays of sarcomeres. The first observed defects in the Tmod-KD myofibers were diagonal myofibrils at the myofiber ends of segments A1 and A2 in second instars (Fig. 2A; Fig. S2). In early third instars, these defects also appeared in mid-segments (A4, A5) and became even more pronounced in the anterior segments, which displayed ectopic diagonally intersecting myofibrils (Fig. 2B,D; Fig. S2). In late third instars, the sarcomeric pattern was disrupted almost entirely on the visceral side of anterior segments and large bundles of F-actin replaced organized myofibrils (Fig. 2C,F; Fig. S2). Occasionally, interlaced “braided” myofibrils formed at the beginning of late third instar, and ectopic radial F-actin accumulations arose by the end of late third instar (Fig. 2E,G). Additionally, once the first diagonally misoriented myofibrils were observed, perpendicular myofibrils were found adjacent to the myotendinous junction (MTJ) at the posterior myofiber end (Fig. 2B). These perpendicular myofibrils also later developed at the anterior end of the myofibers (Fig. 2C). Nuclear irregularities appeared in anterior segments during late third instar stage and, together with ectopic F-actin structures, were only observed in anterior segments. These data indicated Tmod-KD myofibers experience a specific temporal and spatial pattern of structural changes, which appeared in parallel with larval growth, with the most severe disruptions occurring in dramatically growing third instar larvae.

**Fig. 2:**
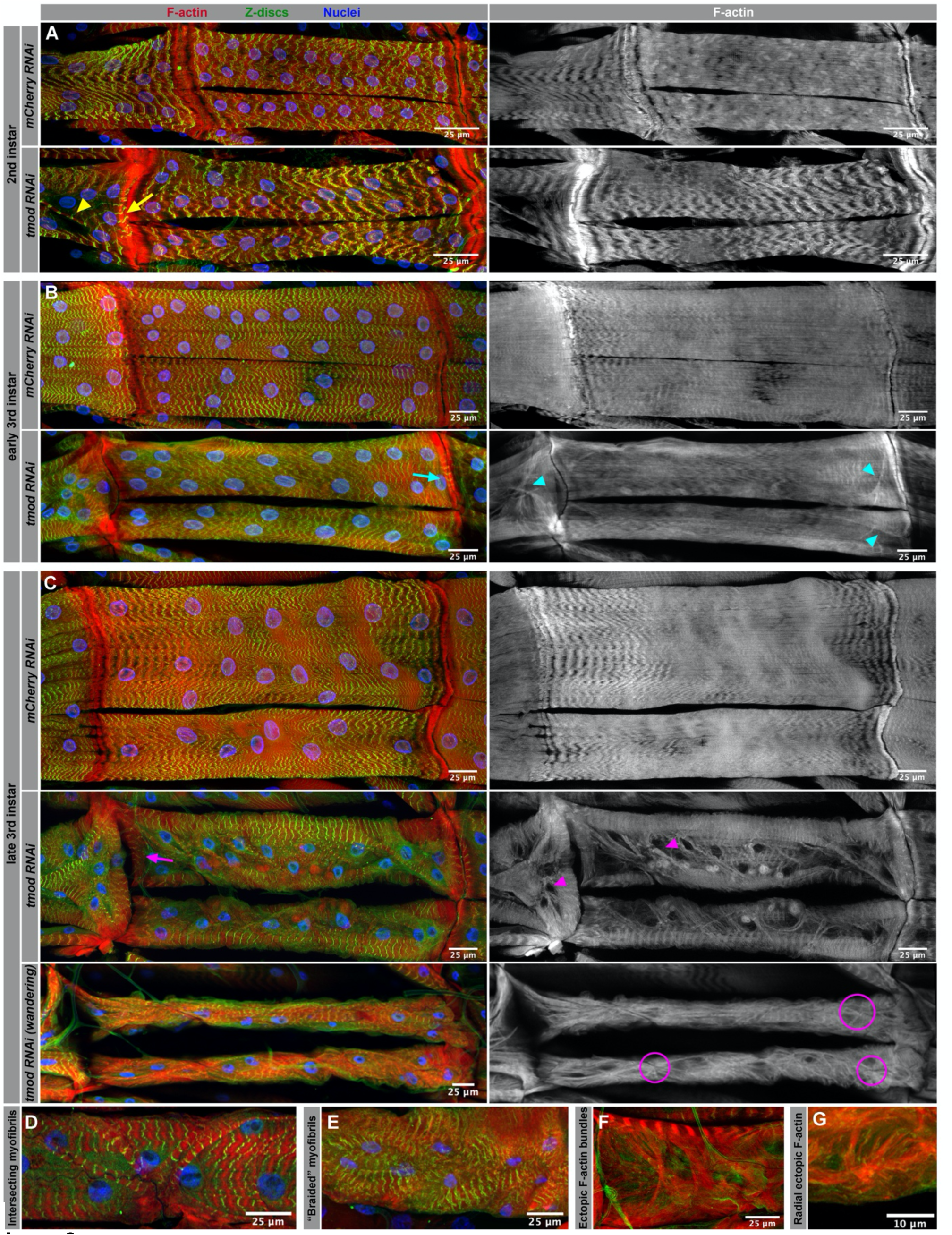
Structural patterned deterioration of Tmod-KD myofibers throughout *Drosophil* larval stages. (A) Posterior end of VI1 myofiber from A1 segment and full length VL3 and VL4 myofibers from A2 segment in control and Tmod-KD second instar larvae (red, phalloidin; green, Zasp::GFP and blue, Hoechst) (left). F-actin channel only (gray, phalloidin) (right). Yellow arrowhead: diagonally misoriented myofibril in VI1 from A1. Yellow arrow: perpendicular misoriented myofibrils in VI1 from A1. (B) Worsening myofibers from A1 and A2 segment in Tmod-KD early third instars (bottom), controls (top). Cyan arrowheads: diagonally misoriented myofibrils. Cyan arrow: perpendicular myofibrils in posterior end of VL3 muscle from A2 (C) Worsening myofibers from A1 and A2 in Tmod-KD late third instars (middle panels) and wandering late third instar (bottom panel), control (top panel). Magenta arrowheads: worsening of diagonally misoriented myofibrils from A1 and A2. Magenta arrow: perpendicular myofibrils in anterior end of VL3 muscle from A2. Magenta circles: ectopic F-actin structures that replace stereotypical myofibrillar pattern. Quantifications across the A-P axis in Supplemental Fig2D. (D) Intersecting grouped myofibrils in early third instar. (E) Interlaced or “braided” myofibrils in late third instar. (F and G) Ectopic F-actin bundles and radial actin structures, respectively, that lack myofibrillar pattern in wandering late third instar. N=3 replicates, each with n=5 larvae per stage Scale bars: 25μm (A-F) and 10μm (G).

### Tmod’s role as a pointed-end capping protein is conserved in *Drosophila*

To understand how Tmod loss generates these phenotypes, we studied the biochemical activity of *Drosophila* Tmod using purified proteins *in vitro*. Tmod’s canonical role in vertebrates is filament pointed-end capping (Dye et al., 1998, Weber et al., 1994, Gregorio et al., 1995, Mardahl-Dumesnil and Fowler, 2001, Fowler and Dominguez, 2017). However, some Tmods also have limited actin filament nucleation activity (Fischer et al., 2006, Yamashiro et al., 2010, Yamashiro et al., 2014), mimicking the activity of Tmod’s structural homolog Lmod that has strong nucleation activity (Chereau et al., 2008). Lmod is not found in *Drosophila*. Nonetheless, while mammals express only four Tmod isoforms, *Drosophila* expresses 18 different splice variants. Moreover, several *Drosophila* Tmod isoforms differ substantially in sequence from their mammalian counterparts, displaying N- and/or C-terminal extensions reminiscent of those present in mammalian Lmods (Fig. S3F). These observations led us to ask whether *Drosophila* Tmod isoforms capped pointed-ends or nucleated filaments.

We chose two isoforms for *in vitro* analysis, the selection of which was based on several criteria: the RNAi target sites, the abundance of the transcript in control muscle, the degree of KD, sequence analysis, and the unique structures of the isoforms (Fig. S3A-F, Table S3). Based on these factors, isoforms Q, O, and K, representing two different N-terminal subgroups, were chosen for in-depth biochemical analysis (note that isoforms O and K have identical protein sequences). We expressed and purified TmodQ, TmodO/K, and human TMOD3 (a control for capping activity) in *E. coli* and monitored the effect of TmodQ and TmodO/K on actin assembly using the pyrene-actin polymerization assay (Doolittle et al., 2013). Because the concentration of actin monomers (1.5 μM) is higher than the critical concentration for monomer addition at the pointed-end (∼0.6 μM), pointed-end capping under these conditions would be observed as a decrease in fluorescence with increasing Tmod concentrations, resulting from inhibition of monomer addition at the pointed-end. In contrast, if Tmod nucleated filaments from monomers we would observe an increase in fluorescence (i.e. polymerization), resulting from the formation and elongation of new barbed-ends. Similar to human TMOD3, TmodQ and TmodO/K reduced polymerization in a concentration-dependent manner (Fig. 3A; Fig. S3G,H). These results suggested that the two representative *Drosophila* Tmod isoforms analyzed function as *bona fide* pointed-end cappers like their mammalian Tmods, and not nucleators like mammalian Lmods. Therefore, the observed KD phenotypes are likely due to a deficiency in pointed-end capping.

**Fig. 3:**
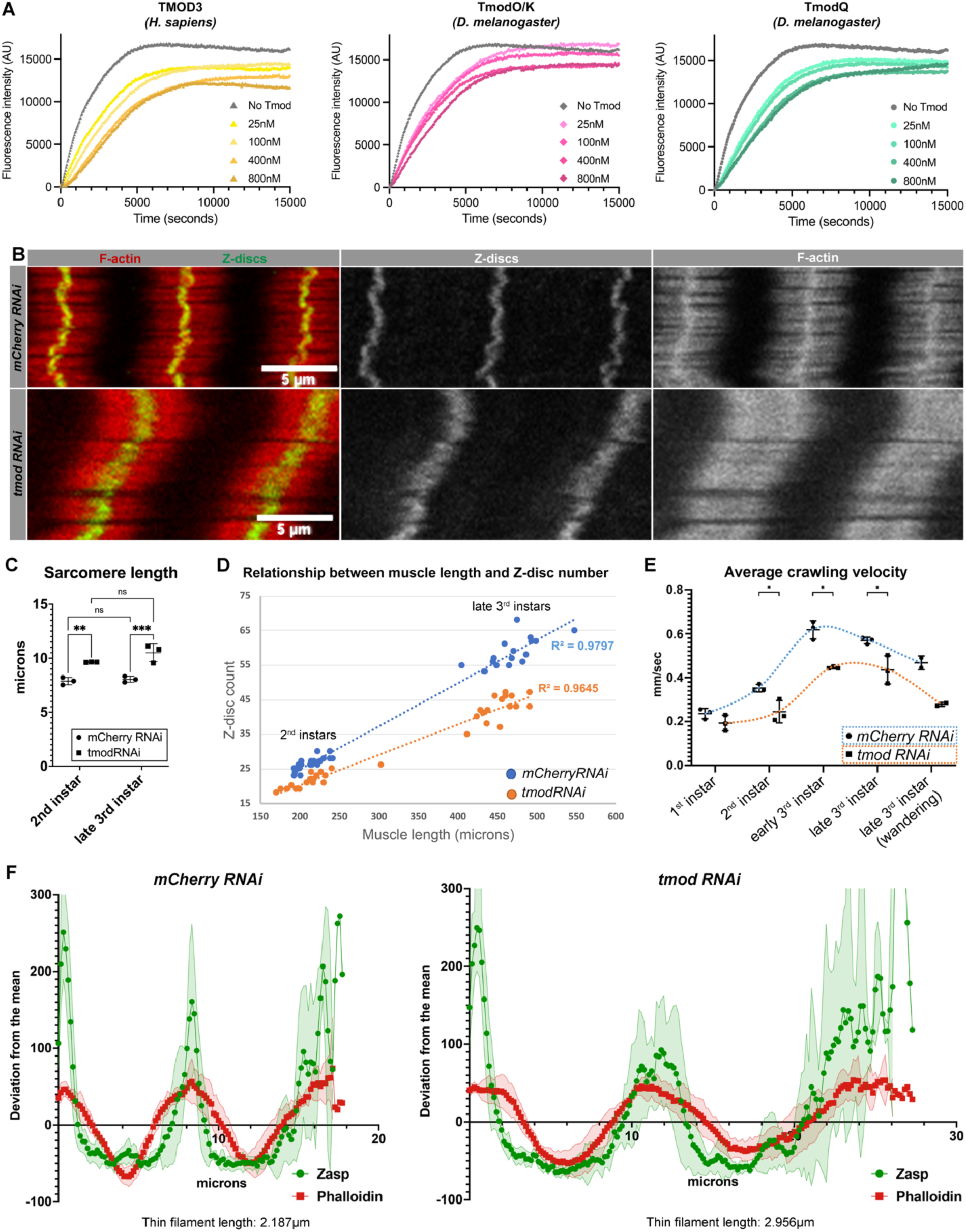
*Drosophila* Tmod is a capping protein and Tmod-KD myofibers display longer thin filaments and sarcomeres. (A) Pyrene-labeled actin polymerization assays at the pointed-end with human TMOD3 *Drosophila* TmodO/K and TmodQ at multiple concentrations (N=4 per isoform per concentration). (B) Cuticular sarcomeres from control and Tmod-KD myofibers of late third instars in A2 (red, phalloidin, and green, Zasp) (left). Middle panel: Zasp channel; right panel: phalloidin channel. (C) Sarcomere length in control and Tmod-KD VL3 myofibers from A2 in second instars and late third instars. Each dot is one experiment (N=3 replicates, each with n_2nd instars_=75-135, n_late 3rd instars_=45-105 sarcomeres [from 3-8 different larvae] per genotype; p_2nd instars_=0.0076, p_3rd instars_=0.0009, pm_CherryRNAi_=0.9703, p_tmodRNAi_=0.1843, ordinary 2-way ANOVA multiple comparisons). Mean ± SD. (D) Scatterplots and regression lines (dotted) comparing Z-disc number in relationship to muscle length (in microns) in control (blue) and Tmod-KD (orange) VL3 myofibers from A2 in second instars and late third instars (N=3 experiments, each with n_2nd instars_=19-20, n_late 3rd instars_=16-17 myofibers per genotype). R² are indicated. (E) Crawling velocities from each larval stage in control (blue) and Tmod-KD (orange). Each dot is one experiment (N=3 replicates, n_1st instars_=10, n_2nd instars_=10, n_early 3rd instars_=5-10, n_late 3rd instars_= 3-10, n_wandering_=3-4 larvae per genotype and experiment; p_1st instars_=0.245, p_2nd instars_=0.0326, p_early 3rd instars_=0.0182, p_late 3rd instars_=0.0433, p_wandering_=0.0507, two-tailed paired Student’s t-test). Mean ± SD. (F) Intensities of phalloidin (red) and Zasp::GFP (green) normalized as the deviation from the mean and plotted across the length of two sarcomeres in A2 of late third instars. Thin filament length is obtained from the length under the center peak divided in half (n=4-6 muscles from different larva per genotype). Mean ± SD. Scale bars: 5μm (B)

### Tmod regulates thin filament and sarcomere length

Previous work has shown that Tmod regulates sarcomere length by capping the pointed-end of sarcomeric F-actin [reviewed in Littlefield and Fowler (2008), Ghosh and Fowler (2021)]. To assess the effects of Tmod-KD in *Drosophila* larval myofibers, we quantified muscle length, Z-disc number, and sarcomere lengths in second and late third instars. At both timepoints, Tmod-KD myofibers had fewer Z-discs than controls with the same length (Fig. 3D; Fig. S4A,B). Z-disc numbers were reduced by 15% in the second instar and by 26% in late third instar, indicating that sarcomere size increased in growing Tmod-KD myofibers. Accordingly, the distance between Z-discs revealed a significant 22% and 31% increase in sarcomere length in Tmod-KD myofibers at the second and late third instar stages, respectively (Fig. 3C). To confirm that Tmod regulates sarcomere size via its effect on thin filaments, we quantified sarcomeric F-actin length in late third instars. Thin filament length increased by 35% in Tmod-KD myofibers. Interestingly, the proportion of thin filament length to sarcomere length was similar to control, whereby thin filaments account for 25% of sarcomere length (Fig. 3B,F). Together, these data suggested that capping of sarcomeric F-actin by Tmod is required to prevent increases in sarcomere size during myofiber growth. The same thin filament to sarcomere length ratio in Tmod-KD and control myofibers indicates that independent mechanisms maintain sarcomere proportions.

Sarcomere size is highly conserved for optimal contraction, and changes in its size have been linked to reduced muscle function (Fernandes and Schock, 2014, Molnar et al., 2014, Spletter et al., 2018, Spletter et al., 2015). To assess the functional consequences of muscle-specific Tmod KD, we performed larval locomotion assays. We analyzed larval velocity throughout larval development and myofiber function significantly decreased, starting at second instar stage and over larval development (Fig. 3E). These data suggested Tmod-KD caused myofiber weakness even before obvious muscle deterioration, and that changes in thin filament length could be the first effect of Tmod KD to reduce muscle function.

### Tmod stabilizes core components of the muscle attachment site

The current model of myofibrillogenesis suggests that tension is crucial for this process. Specifically, the muscle attachment sites at the MTJs play a fundamental role in generating tension within a myofiber. The MTJ represents an analogous structure to focal adhesion sites (Atherton et al., 2016, Martino et al., 2018, Pardo et al., 1983, Quach and Rando, 2006, Mohammad et al., 2012), which are composed of the Vinculin-Talin-Integrin complex tethering the actin cytoskeleton to the ECM, or in the case of the myofiber, the myofibrils to the muscle attachment site. We hypothesized that Tmod functions at the MTJs to stabilize myofibril attachment during myofiber growth.

In agreement with this hypothesis, we detected Tmod localized at the ends of control myofibers, adjacent to myofibrillar F-actin and co-localizing with β_PS_-Integrin (Mys) (Fig. 4A,C). We next evaluated protein enrichment of the tension-mediating proteins β_PS_-Integrin and its binding partner Talin (Rhea) in Tmod-KD MTJs. β_PS_-Integrin and Talin were significantly decreased in Tmod-KD anterior segments (Fig. 4E,F; Fig. S5A-C), however we did not detect myofiber detachment. To determine whether the strong decrease in signal was due to mislocalization or an overall decrease in protein levels, we performed Western blot analysis. Talin levels did not decrease in Tmod-KD wandering larvae muscle-enriched lysates (Fig. 4B), suggesting that Talin is mislocalized. Upon further examination, we observed pools of β_PS_-Integrin and Talin in ectopic areas (Fig. 4F). These data suggested Tmod’s F-actin capping function is required for establishing the proper anchorage sites at the myofiber ends by stabilizing the interaction between F-actin and tension-mediating proteins.

**Fig. 4:**
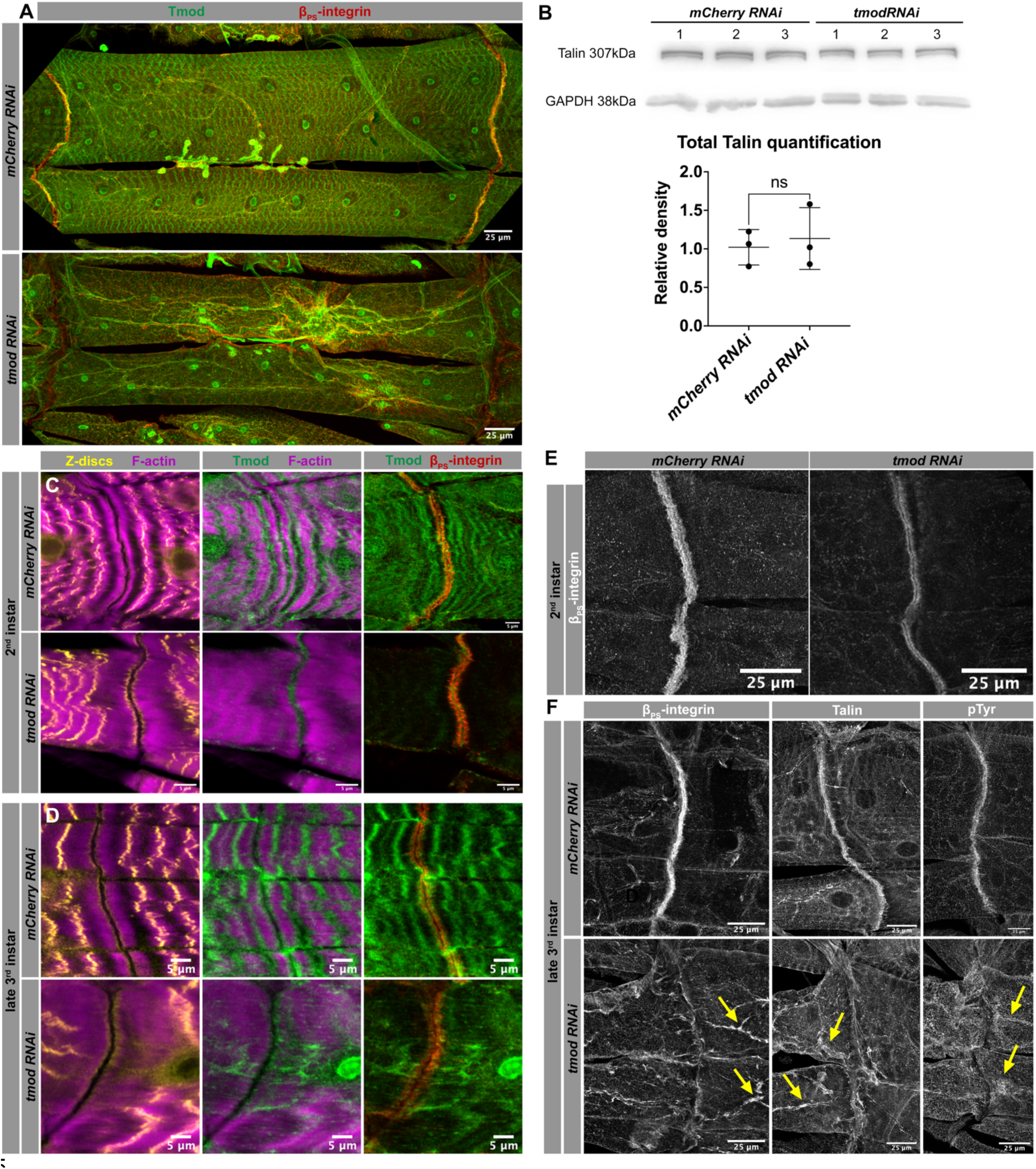
Tmod-KD muscles exhibit Tmod decrease and loss of mechanosensory protein enrichment and tension at the MTJ. (A) VL3 and VL4 myofibers in A2 from late third instars in control and Tmod-KD larvae. In control, Tmod (green) displays a stereotypical pattern in the sarcomeres and its colocalization with β_PS_-integrin (red) at the MTJ (top panel). In Tmod-KD myofibers, Tmod is not present at the same intensity levels at the sarcomeres nor the MTJ (bottom panel). (B) Talin Western blot and quantification in late third instar muscle-enriched lysates. Glyceraldehyde 3-phosphate dehydrogenase (GAPDH), loading control. Each dot is an experiment (N=3 replicates, each with n=5-10 larval carcasses per genotype, p=0.7846, two-tailed paired Student’s t-test). Mean ± SD. (C and D) Analysis of protein localization (magenta, phalloidin; yellow, Zasp::GFP; green, Tmod; and red, β_PS_-integrin) relative to one another at the MTJ (A2-3) in control and Tmod-KD in second and late third instar larvae. (E) β_PS_-integrin (white) strong enrichment at the MTJ (A2-3) in second instar control compared to decreased signal in Tmod-KD. Quantification in Supplemental Fig 5A. (F) β_PS_-integrin (left), Talin (white, middle) and pTyr (white, right) strong enrichment at the MTJ (A2-3) in wandering third instar control compaed to absence of signal in Tmod-KD. Quantifications in Supplemental Fig 5B. Yellow arrows: ectopic pools of protein. Scale bars: 5μm (C, D) and 25μm (A, E, and F).

To address whether mislocalization of β_PS_-Integrin and Talin correlated with tension loss at the MTJ, we assessed phospho-tyrosine (pTyr), a marker of active cytoskeletal tension sites (Chrzanowska-Wodnicka and Burridge, 1994, Martino et al., 2018, Parsons et al., 2010). pTyr was significantly decreased in Tmod-KD MTJs (Fig. 4F; Fig. S5D), indicating reduced tension. Together, these experiments suggested a decrease in mechansensory proteins and a reduction of tension at the muscle ends. A lack of tension could cause myofibrillar disorganization and impaired myofibrillogenesis in the Tmod-KD myofibers.

### Tmod regulates myonuclear morphology, positioning, and function

Cytoplasmic F-actin is found in and around the nucleus (Davidson and Cadot, 2021, Caridi et al., 2019). These filaments play roles in gene expression, in the cell’s response to mechanical stimuli, and in nuclear positioning in several systems (Farina et al., 2016, Verheyen and Cooley, 1994, Chang et al., 2015). We hypothesized Tmod-KD could affect cytoplasmic F-actin to generate the nuclear phenotypes that we observed. Control *Drosophila* larvae displayed multiple, evenly spaced, round disc-shaped nuclei on the myofiber’s visceral side. In contrast, many Tmod-KD nuclei were elongated, with an increased perimeter compared to control (Fig. 5A-C). Along the AP axis, the most irregular-shaped nuclei were located towards the middle of the myofiber (Fig. 5D). It is possible Tmod-KD directly affected F-actin dynamics that stabilize nuclear shape and position. Unfortunately, imaging of cytoplasmic and/or nuclear F-actin is obstructed by the bright signal of the underlying sarcomeres, which prevented the further analysis of direct effects of Tmod-KD on cytoskeleton and nucleoskeleton in this study.

**Fig. 5:**
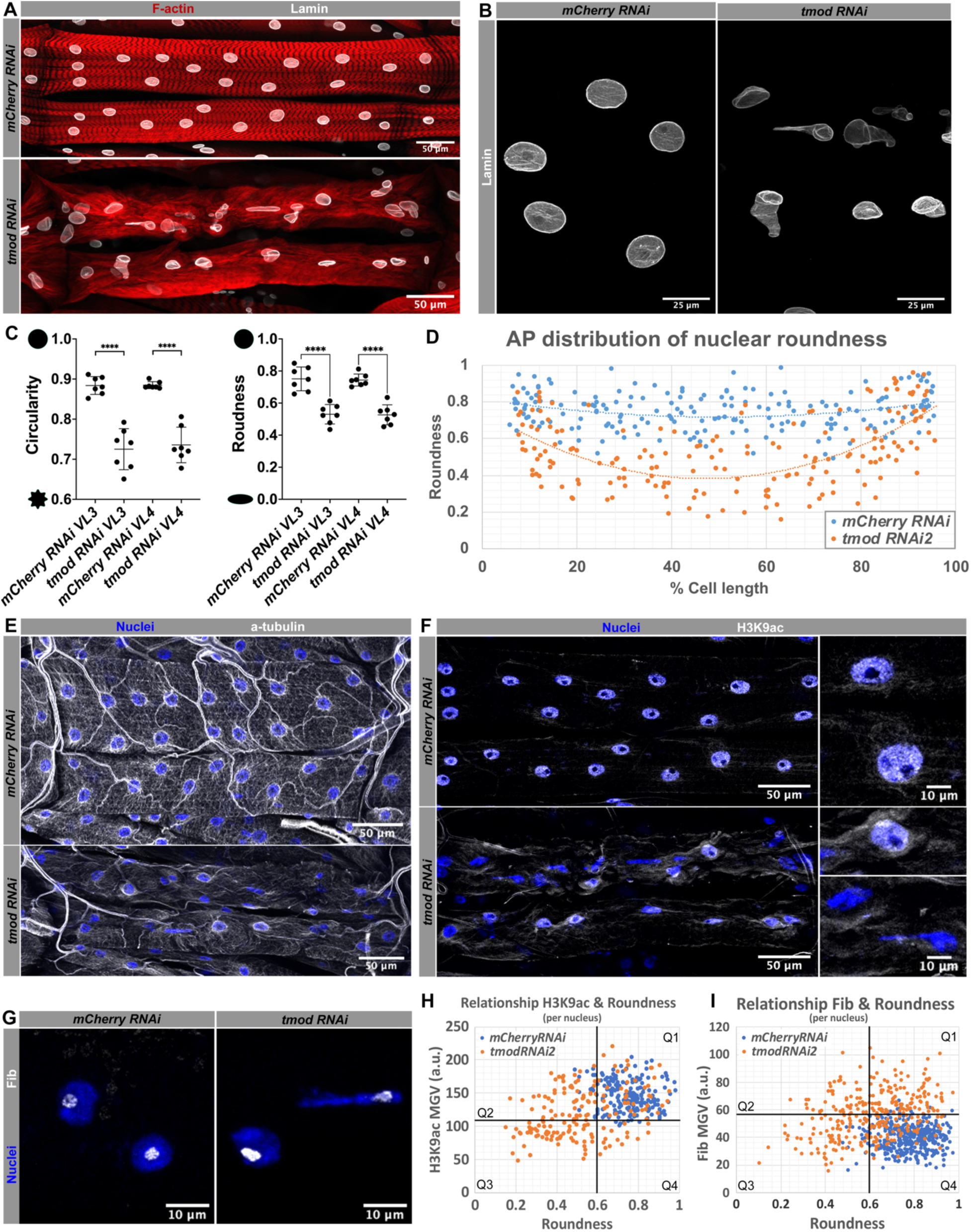
Tmod-KD myofibers display misshapen and dysfunctional nuclei in late third instars. (A) VL3 and VL4 myofibers (red, phalloidin) with nuclei (white, Lamin) in A2 in control (top) and Tmod-KD larvae (bottom). (B) High magnification of control myonuclei (white, Lamin) and misshapen and internalized Tmod-KD nuclei. See supplemental Movie 1 and 2. (C) Quantification of two shape descriptors: circularity and roundness in VL3 and VL4 myofibers in A2 from control and Tmod-KD. Graph represents one experiment, each dot is one myofiber per larva (n_VL3_= 15-20, n_VL4_=8-11 myonuclei per myofiber and genotype; circularity p_VL3_<0.0001, circularity p_VL4_<0.0001; roundness p_VL3_<0.0001, roundness p_VL4_<0.0001, ordinary one-way ANOVA multiple comparisons). Experiment repeated N>3. Mean±SD. (D) Distribution of nuclear roundness along the AP axis of the myofiber from control (blue) and Tmod-KD (orange) myofibers. Polynomial trendline shown by curved dotted lines. (E) MT staining (white, α-tubulin; blue, nuclei) showing radial perinuclear arrays and longitudinal tracks in control and a decrease of these in Tmod-KD. (F-G) H3K9ac and Fib staining (white) indicating active transcription sites and nucleolus respectively in myonuclei both in control and Tmod-KD myofibers. (H and I) Scatterplots showing H3K9ac and Fib pixel intensity levels (mean gray value) in relationship to roundness levels. Each graph represents one experiment, each dot is one myonucleus (n_H3K9ac_=178-190 myonuclei per genotype, from 7 VL3 and VL4 muscles sets, in 7 larvae; n_Fib_=308-396 myonuclei per genotype, from 12-15 VL3 and VL4 muscle sets, in 7-8 larvae). (H) Q3 indicates Tmod-KD misshaped nuclei (roundness<0.6) with decreased H3K9ac intensity when compared to control in Q1. (I) Q1 indicates Tmod-KD WT-like nuclei (roundness>0.6) with increased Fib intensity when compared to control in Q4. Q3 indicate Tmod-KD misshaped nuclei (roundness<0.6) with a similar Fib intensity when compared to control Experiments repeated N>2 unless specified otherwise. Scale bars: 50μm (A, E and F), 25μm (B) and 10μm (F and G).

Stable radial microtubule (MT) arrays around the nuclei maintain positioning and nuclear integrity during myofiber contraction (Zheng et al., 2020, Becker et al., 2020, Manhart et al., 2018, Metzger et al., 2012). In addition, MTs physically interact with actin (Farina et al., 2016, Dogterom and Koenderink, 2019), which raises the possibility that actin-MT crosstalk supports nuclear shape and position. To assess if perinuclear MTs were affected in Tmod-KD muscles, we quantified the number of intact radial MT arrays in control and Tmod-KD myofibers. We observed a significant decrease in intact perinuclear MT arrays in Tmod-KD muscles and noted a decrease in longitudinal MT-tracks (Fig. 5E; Fig. S6A). These data indicated that Tmod affects critical MT networks in myofibers, by mediating the interactions of the MT cytoskeleton with actin filaments in/around the nuclei. A loss of perinuclear MTs could subsequently result in destabilization of nuclear shape and position.

To test if nuclear activity was affected by nuclear morphology, we analyzed the transcriptional and translational potential of Tmod-KD myonuclei. To assess transcription, we quantified H3K9ac, a marker of open chromatin and transcriptionally active regions in the DNA (Kharchenko et al., 2011). Nuclei with the most irregular shapes (roundness<0.6) exhibited decreased H3K9ac levels, whereas WT-shaped nuclei (roundness>0.6) showed similar H3K9ac levels to control (Fig. 5F,H; Fig. S6B). To assess translation, we quantified nuclear intensity of Fibrillarin (Fib), a nucleolar protein that plays an essential role in ribosome biogenesis and is a commonly used readout for protein synthetic ability (Gillery et al., 1996, Lefevre et al., 2001). Misshaped nuclei (roundness<0.6) exhibited similar Fib intensity levels to control, while WT-shaped nuclei (roundness>0.6) showed higher levels than control (Fig. 5G,I; Fig. S6C). These data suggested that loss of nuclear shape and position in Tmod-KD myofibers reduced transcriptional output but increased translational potential. While deformed nuclei are functionally compromised, WT-shaped nuclei in the Tmod-KD muscle appeared to compensate for this deficiency by increasing translation.

### Tmod knockdown alters actin isoforms, degradation, and oxidative metabolism

We analyzed the transcriptome of Tmod-KD larvae to better understand the mechanisms behind the observed phenotypes and uncover additional consequences. Consistent with the morphological features found in Tmod-KD muscle, we detected misregulation of actin genes. Sarcomeric actin isoforms (Act57B, Act87B) (Roper et al., 2005, Fyrberg et al., 1983) were upregulated (Fig. 6; Table S4), consistent with the observed ectopic F-actin bundles (Fig. 2F-G; Fig. 6E). In contrast, cytoplasmic actin isoforms (Act5C, Act42A) were downregulated in Tmod-KD myofibers (Fig. 6E; Table S4), consistent with the alterations in nuclear shape detected in Tmod-KD.

**Fig. 6:**
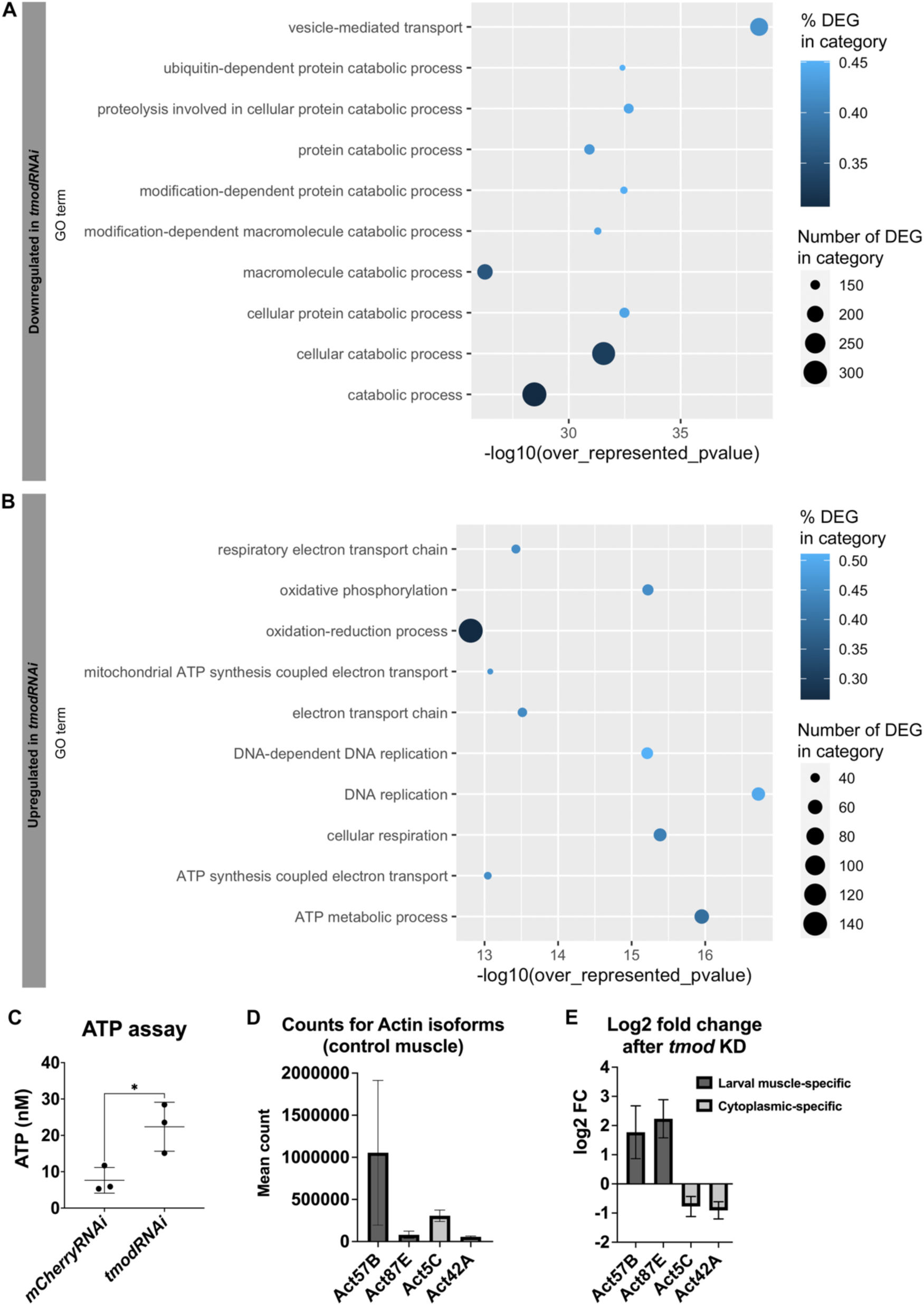
Tmod-KD late third instar larvae show differential gene expression (DEG). (A and B) Top 10 downregulated and upregulated Gene Ontology gene sets in Tmod-KD vs control by ORA. (C) ATP concentration in muscle-enriched larval lysates. Each dot is one experiment (N=3 replicates, each with n=5-10 larval carcasses per genotype, p=0.0285, two-tailed unpaired Student’s t-test). Mean ± SD. (D) Normalized counts for the different *Drosophila* actin genes in control larvae. Mean ± SD. (E) Log2 fold change of actin genes isoforms in Tmod-KD vs control (p_Act57B_=0.0053, p_Act87E_<0.0001, p_Act5E_= 0.0011, p_Act42A_<0.0001). Mean±SE. RNAseq data was used for plots A, B, D, and E (N=3 replicates, each with n=7-10 larval carcasses per genotype).

We also performed an over-representation analysis (ORA) to determine if a set of shared genes was differentially expressed based on Gene Onthology (GO) and Kyoto Encyclopedia of Genes and Genomes (KEGG) gene sets. These data indicated that degradation-related genes (particularly proteasomal genes) and/or pathways were downregulated, while metabolism-linked genes (particularly oxidative metabolism genes linked to mitochondrial ATP production) and/or pathways were upregulated in Tmod-KD muscles (Fig. 6A,B). To test whether ATP production was indeed altered in Tmod-KD muscles, we performed a functional ATPase assay. Consistent with our RNAseq data, ATP levels significantly increased in Tmod-KD myofibers (Fig. 6C). Together, these data suggested a role for Tmod, either directly or indirectly, in gene expression. We speculate that these changes reflect a compensatory mechanism to promote growth in a degenerating muscle.

## DISCUSSION

Tmod function has been studied using various cell types (Fowler, 1997). Here we investigated the effects of Tmod KD in *Drosophila* larval myofibers during development using a combination of detailed *in vivo* analyses, *in vitro* experiments, and transcriptomic approaches. Our studies suggest that through its capping function, Tmod affects actin dynamics in different locations within the myofiber and is ultimately essential for muscle function.

We propose the following model for Tmod’s pleiotropic effects in muscle cells (Fig. 7). In the sarcomere, the barbed end of F-actin is capped by CapZ and anchored to the Z-disc, while the pointed-end is capped by Tmod. However, due to Tmod’s low affinity for F-actin (Weber et al., 1994), there are yet opportunities for monomer addition and therefore, the pointed-end remains suitable for sarcomere growth. When Tmod is knocked-down, thin filaments grow in length. We suggest this additional polymerization causes sufficient imbalance to favor the production of muscle-specific larval G-actin isoforms (Act57B, Act 87E); this increased G-actin availability and continuously growing sarcomeric F-actin would result in ectopic F-actin bundles and radial structures. In the cytoplasm, F-actin growth occurs through the canonical pathway, which is, generally, by the addition of G-actin monomers at the barbed end and depolymerization at the pointed-end. When Tmod is reduced, excessive depolymerization occurs, due to loss of Tmod’s capping activity. This excess of G-actin monomers decreases the production of cytoplasmic-specific actin isoforms (Act5C, Act42A). Loss of cytoplasmic perinuclear and nuclear F-actin would contribute to the loss of nuclear organization and directly or indirectly affect nuclear output. At the MTJ, myofibrils are anchored to the muscle ends via cytoplasmic F-actin. In the absence of Tmod, lack of stable attachments via F-actin impairs tension along the myofibrils, leading to myofibril disorganization and deterioration, and defects in myofibrillogenesis. Protein degradation and metabolism would be affected similarly by destabilization of required structures for myofiber organization or via transcriptional changes. Together, loss of Tmod at these different sites leads to impaired muscle function.

**Fig. 7:**
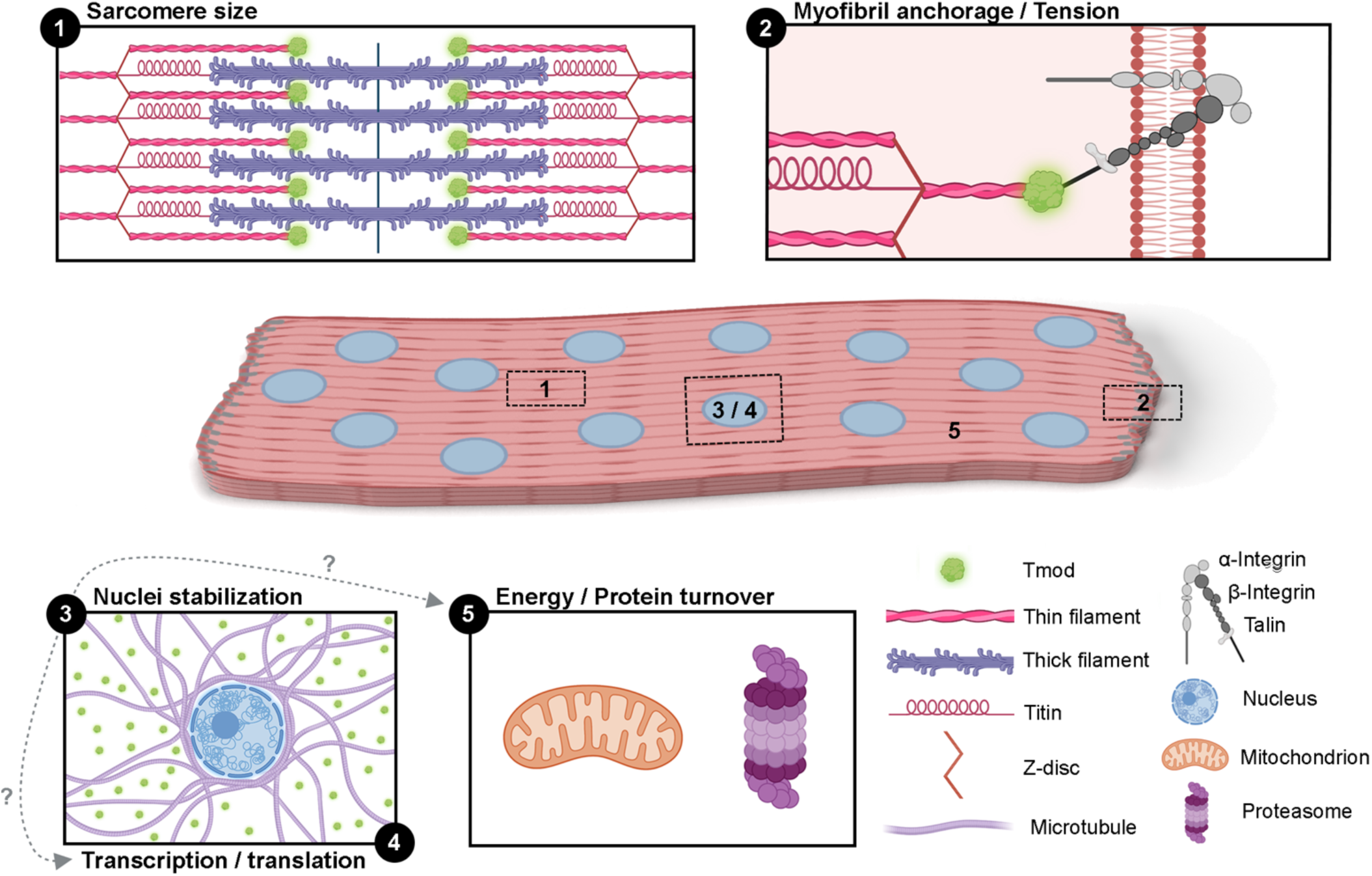
Summary of Tmod’s diverse roles in *Drosophila* larva myofibers. (1) Sarcomere size: Tmod regulates sarcomere and thin filament length. (2) Myofibril anchorage: Tmod establishes tension along the longitudinal axis by stabilizing the interaction of β_PS_-integrin/Talin at the MTJ. (3) Nuclear stabilization: Tmod maintains nuclear integrity either through MT-actin interactions or through nuclear F-actin. (4) Transcription/translation: Tmod regulates nuclear output, through actin as a transcription factor in the nucleus or by actin’s role in nuclear architecture and chromatin organization. (5) Energy/Protein turnover: Upon Tmod-KD, we observed an increase in energy production genes and a decrease of proteasome genes. This change in transcriptome could be due to Tmod’s role in nuclear stabilization or as compensation to generate myofiber growth.Grey dashed arrows and question marks indicate potential consequences.

While every myofiber experiences the same Tmod-KD effect and follows the same pattern of deterioration, not every myofiber is affected equally. Along the larval AP axis, we see that the *tmod* phenotypes appear first and are most severe in anterior abdominal segments which contain larger muscles than posterior segments, suggesting that growth and/or myofiber size may be a contributing factor. However, we observe that some muscles at the same axis position are preferentially affected, which suggests a more complex situation. Freely moving *Drosophila* larvae constantly pivot and sweep their heads, which are supported by muscles of segments A1-A3 (Lahiri et al., 2011); hence, muscle usage may affect muscle deterioration in our model. In parallel, the timing and distribution of muscle pathology in many human muscle diseases, as well as animal models, follow specific patterns (Barohn et al., 2014, Sewry et al., 2019). Studies in vertebrate models suggest that muscle fiber type might play a role [reviewed in Talbot and Maves (2016)]. The factor(s) that make specific muscles more susceptible to deterioration remain unknown. The *Drosophila* larval musculature, consisting only of one fiber type, provides a useful system for further studies in this context.

Alternative splicing results in a total of 18 Tmod isoforms. Why the fly produces so many different isoforms is unclear. We show that isoforms Q and O/K display Tmod’s characteristic F-actin capping function, despite having N-terminal extensions comparable to human LMOD. We did not find biochemical evidence of F-actin nucleation. It is possible that one of the remaining Tmod isoforms (most notably the isoforms with C-terminal proline-rich extensions) may possess nucleation activity like that of human LMODs. However, the extremely low levels of such isoforms in our RNAseq data suggest they are not expressed in myofibers, or at the tested developmental stages.

Tmod is suggested to define thin filament and sarcomere length in a variety of model systems (Szikora et al., 2022). In adult *Drosophila*, Tmod overexpression results in shorter sarcomeres and thin filaments (Molnar et al., 2014, Mardahl-Dumesnil and Fowler, 2001). In vertebrate systems, such as mouse Tmod-null mutants, thin filament length increases (Littlefield and Fowler, 2008). *In vitro* rat cardiomyocytes show increased or decreased Tmod levels resulting in shorter or longer sarcomeric actin filaments, respectively (Sussman et al., 1998). Our data demonstrate that Tmod-KD in *Drosophila* larval myofibers increases thin filament and sarcomere length. Thin filament to sarcomere length ratio was maintained in Tmod-KD myofibers. This ratio was similar to measurements in frog sartorius (26%) and human pectorialis (30%) (Gokhin and Fowler, 2013). This indicates that mechanisms that regulate sarcomere proportions are conserved and regulated independently of Tmod function. We further find sarcomere size increases throughout larval growth, indicating that the uncapped pointed-ends of sarcomeric actin filaments do not respect a specified size, exhibiting continuous polymerization in Tmod’s absence. In healthy muscle, sarcomere length is fixed to optimize muscle contraction, forcing growing cells to add sarcomeres rather than increasing their length (Balakrishnan et al., 2020, Williams and Goldspink, 1971, Dix and Eisenberg, 1990, Goldspink, 1983, Haas, 1950). We show an inverse relationship between sarcomere number and length in Tmod-KD myofibers, suggesting that myofibrils could grow by increasing their length via sarcomere growth. Consistent with recently published work, our data indicates that muscle length and sarcomere number are regulated independently of one another (Brooks et al., 2022). Loss of this adaptive behavior is associated with myopathy progression (Ottenheijm et al., 2009, Winter et al., 2016). Moreover, the decrease in myofiber size is suggestive of compromised myofibril addition to increase muscle mass, which further supports that mechanisms of muscle growth (addition of new sarcomeres and hypertrophy) are affected in Tmod-KD larvae. Our analyses show locomotive defects (first instar) before discernable changes in myofiber structure (second instar). While technical limitations prevented the detailed analysis of first instar sarcomeres, we suggest that changes in thin filament length might be the first effects of Tmod KD that directly affect muscle function. In parallel with disruption of sarcomeres and myofibrils, Tmod-KD myofibers display accumulation of ectopic and disorganized F-actin. The location of these ectopic bundled and radial F-actin accumulations near disorganized myofibrils indicates that continuously growing thin filaments could contribute to these ectopic structures. Whether the ectopic disorganized F-actin filaments were intended to serve as a myofibril template (Fenix et al., 2018, Dlugosz et al., 1984) or they result from deteriorating myofibrils remains unclear.

The currently favored hypothesis in the field describes sarcomere and myofibril formation as a tension-driven self-organizing process, whereby tension generated through muscle attachment at the MTJs is used to orient and longitudinally assemble sarcomeres and myofibrils (Lemke and Schnorrer, 2017). Here, we show that Tmod-KD results in mislocalization of two core mechanosensitive proteins and a reduction in tension at the muscle ends. According to the hypothesis, lack of tension at the muscle ends affects addition of new sarcomeres as well as homeostasis along the entire myofibril. This is consistent with our Tmod*-*KD data showing reduced sarcomere number and aberrant myofibril organization and stability. Previous studies using *tmod* mutant zebrafish also reported defects in myofibril organization and muscle weakness (Berger et al., 2022, Berger et al., 2014), but a role for Tmod in sarcomere addition had not been described. Interestingly, we observe perpendicular myofibrils exclusively localized at the myofiber ends, which demonstrate sarcomeric organization albeit incorrect orientation. This raises the possibility that due to the lack of longitudinal stability in this growth zone, the forming sarcomeres and growing myofibrils orient towards the next best structure that provides resistance, connecting to the MTJ, existing myofibrils or the visceral plasma membrane. A similar phenotype with misoriented myofibrils was observed in *Drosophila* larval muscle expressing the constitutively active form of C3G, a Rap1 activator (Shirinian et al., 2010). Importantly, Rap1 has been demonstrated to recruit Talin to the cell membrane (Shattil et al., 2010, Lee et al., 2009). We find ectopic localization of mechanosensitive proteins along the plasma membrane of Tmod-KD myofibers, which could lead to ectopic F-actin accumulations. This could result in local changes in tension throughout the myofiber and contribute to myofibril misalignment. Together, our study suggests that Tmod is crucial for maintaining alignment, stability, and growth of myofibrils. Our data are consistent with the tension-driven self-organizing myofibrillogenesis hypothesis and adds a role for Tmod in tension mediation.

Our data implicates Tmod in nuclear positioning in the Z-axis and maintaining nuclear morphology. Mispositioned and deformed nuclei are common in many myopathies and laminopathies (Romero, 2010, Steele-Stallard et al., 2018, Ross et al., 2019), highlighting the relevance of correct nuclear positioning for proper muscle function. Actin plays a structural role in the nucleus (Bettinger et al., 2004); hence Tmod, through its capping function, may stabilize nuclear and/or perinuclear actin to directly support nuclear integrity. We did not detect nuclear or perinuclear actin filaments, possibly due to the strong F-actin signal of the adjacent thin filaments. However, we found defects in the MT networks surrounding the nuclei of Tmod-KD myofibers. MTs interact with actin filaments and are crucial in myonuclear positioning and integrity. Thus, Tmod-KD likely has indirect effects on myonuclei by disrupting actin-MT interactions. In addition, defects in nuclear positioning and morphology could be indirectly affected by the loss myofibril structure and orientation. In Tmod-KD myofibers, nuclei are initially located at the periphery, but subsequently sink as myofibril structure is lost. Force exertion on the internalized nuclei could then lead to deformation of nuclear shape. These nuclei also show defects in transcriptional activity. Interestingly, WT-shaped nuclei in Tmod-KD myofibers display increased translation makers, consistent with a compensatory mechanism. In addition to the effects of nuclear morphology, it is possible that Tmod regulates transcription and/or translation via its effect on actin dynamics. G-actin has many important functions in the nucleus and nucleolus and directly regulates gene transcription (Visa and Percipalle, 2010, Bettinger et al., 2004). More research is needed to determine how deviation from normal positioning in 3 dimensions (X,Y,Z) results in impaired muscle function. To this end, future studies focused on the distribution of gene products throughout the myofiber when nuclear positioning is aberrant would be informative.

Our transcriptome analyses suggest that Tmod KD affects two main processes, proteasomal degradation and mitochondrial metabolism, both critical for myofiber growth (Schiaffino et al. 2013). However, the decrease in degradation and increase in mitochondrial metabolism detected in Tmod-KD myofibrils is inconsistent with the observed impaired muscle growth and muscle weakness. We suggest that this transcriptomic profile indicates a compensatory attempt of the myofibers to grow, counteracting the lack of tension along the myofiber and structural deterioration. It is possible that actin inside the nucleus causes these changes through structural and/or transcriptional changes (Chase et al., 2013, Visa and Percipalle, 2010). However, similar myofiber phenotypes and changes in metabolism have been observed in *Drosophila TRIM32* (*thin*) mutant larval muscles (LaBeau-DiMenna et al., 2012, Bawa et al., 2020). Loss of *Drosophila* TRIM32, an E3-ubiquitin ligase, resulted in a decrease in glycolytic intermediates, indicating that TRIM32 switches metabolism from oxidative to glycolytic. The mechanisms by which TRIM32 and Tmod are connected remain to be elucidated, yet the similarities in phenotype and transcriptome suggest they might be interacting genetically or biochemically. This would indicate a role for Tmod in dampening oxidative metabolism and, therefore, ATP production. Strikingly, structural components involved in these pathways, such as mitochondria, are located in high densities at the visceral muscle surface and around myofibrils; hence, they could be affected indirectly via the destabilization of the underlying myofibrils.

Myopathies are often devastating disorders characterized by impaired muscle function that are frequently associated with sarcomeric mutations. The molecular details underpinning this relationship are poorly understood. Our study shows a series of phenotypes associated with KD of one critical thin filament component, Tmod, that reach far beyond the sarcomere. Our findings have important implications, as they suggest a key role for Tmod in several cellular processes, including the regulation of actin isoform expression, the stabilization of tension-mediating proteins, and nuclear integrity and positioning, therefore providing additional insight to the pathogenesis of myopathies. The mechanisms directly linking Tmod and critical myofiber processes, including proteasomal degradation and oxidative metabolism, remain to be fleshed out. Furthermore, our findings support and strengthen the tension-driven self-organizing myofibrillogenesis model. Taken together, this study adds significantly to the understanding of Tmod in muscle development, maintenance, and disease, making it a potential target for new therapeutics that address muscle weakness.

## METHODS

### *Drosophila* husbandry, stocks and crosses

*Drosophila* stocks and experimental crosses were grown on standard cornmeal medium at 25°C in 12:12 Light:Dark conditions under humidity control. The Gal4-UAS system (Brand and Perrimon, 1993) was used for RNA interference studies. Details on the screen that lead to our analysis of *tmod* are found in Table S1 and S2. For muscle-specific expression and RNAi potentiation we tested different driver lines (*UAS-dicer2;;Mhc-Gal4,Zasp66::GFP*, *UAS-dicer2;;Dmef2-Gal4,Zasp66::GFP/Tm6,Hu,Tb* and *UAS-dicer2;;Dmef2-Gal4/Tm6,Hu,Tb*) in combination with *3 UAS-tmod-RNAi* lines (BDSC#41718, BDSC #31534 and VDRC #108389) to validate on-target effects and explore KD levels. The driver lines were crossed with *UAS-mCherryRNAi* (BDSC #35785) as the *control RNAi.* In all instances, larvae displaying a Tb phenotype (indicatory of balancer chromosome) were excluded from the F1 analyzed. The mutant stock *tmod^MI02468^*(BDSC #36446) was also used for phenotype validation with *yw* as control. Except where specified, we used a Dmef2-Gal4 driver and one *UAS-tmodRNAi* line (BDSC#41718) for analyzing Tmod-KD phenotypes (the line is referred to as “*tmod”*). Table S5 contains details on fly stocks.

### Larval staging

For all larval experiments, larvae were kept at 25°C and both male and female larvae were analyzed. Staging was performed on apple juice agar plates, either in the embryo (see Viability assay), or by selecting larvae hatched within a 2h period. First instars intended for experiments were maintained in an apple juice agar plate with yeast paste for ease of manipulation. Larvae for experiments at later stages (locomotion, muscle analysis) were transferred to standard food vials. Larvae were removed from food vials using 30% sucrose solution at second instar stage (∼24h after transfer), early third instar (∼48h after transfer), late third instar (∼96h after transfer) and wandering (∼118h after transfer) stage. Myofiber structure time-course analysis was performed on all four distinct larval stages, in VI1 or VL3/VL4 muscles of anterior segments (A1 and A2). For the RNAseq experiment, late third instars (∼90h after transfer) were processed for dissection. For all other experiments at late third instar stage (∼3.5 days), staging was confirmed by using developmental landmarks such as mouth hooks, and spiracle morphologies (Bodenstein, 1950).

### qRT-PCR

Ten third instar wandering larvae were dissected and rinsed in ice cold HL3.1 medium (Feng et al., 2004) as described in (Brent et al., 2009). Total RNA was extracted from the muscle enriched larval fillets using TRIzol reagent (ThermoFisher #15596026), followed by cleanup with TURBO DNA-free Kit (Ambion #AM1907). The SuperScript III First-Strand Synthesis System for RT-PCR (Invitrogen #18080-051) was used to synthesize cDNA and qRT-PCR reactions were performed on the BioRad CFX96 Real-Time PCR system using the SYBR Select Master Mix for CFX (Applied Biosystems #4472937). Three independent mRNA preparations per genotype were collected and were run in triplicate. Fold changes were calculated based on values obtained using the delta-delta Ct method (Livak and Schmittgen, 2001, Schmittgen and Livak, 2008). Product size and uniformity was assessed via melt curve analysis. *Rpl32* was used as a normalization control. Table S3 contains details on the oligonucleotides used.

### Western Blot

Seven to ten late third instar wandering larvae were dissected (See Immunolabeling and Imaging) and transferred into larval lysis buffer (50mM HEPES pH 7.5, 150mM NaCl, 0.5% NP40, 0.1% SDS, 2mM DTT) supplemented with Complete mini protease inhibitor cocktail [one tablet in 10mL of lysis buffer (Roche #11836153001)]. Lysates were homogenized and protein concentrations were determined using the Bradford assay. Equal amounts of protein (20 μg) were run on a 5% (Talin) or 10% (Tmod) polyacrylamide gel and then transferred onto a nitrocellulose membrane (ThermoScientific #88018). Membranes were blocked with either 5% milk or 5% BSA in TBST (Tris-Buffered Saline +0.1% Tween) for 1h at room temperature. Table S5 contains details on primary and secondary antibodies. Immunoreactions were visualized in a KwikQuant Imager (Kindle Biosciences, LLC #D1001) using 1-Shot Digital-ECL (Kindle Biosciences, LLC #R1003) and intensities were quantified using Fiji (NIH). Protein expression was normalized to GAPDH (loading control) within each sample. Figures and quantifications are representative of 3 experiments.

### Larval tracking

Larvae were staged and extracted from food vials (see Larval staging) at first instar, second instar, early third instar, and late third instar. One larva at a time was placed at the center of an apple juice agar plate (8.5cm diameter) and recorded for 45 seconds (Samsung Galaxy S8+). Movies were converted into image sequences (1 image per second) and each larva was tracked using the Manual Tracking plug-in on Fiji. Average velocity was calculated as in (Balakrishnan et al., 2021). For each genotype, we performed three replicates with ∼10 larvae each. Larval length was equivalent in both phenotypes for each developmental stage.

### Plasmids

The cDNA for TmodO/K (NCBI Reference Sequences: NP_001247373.1 / NP_001189319.1) was acquired from DNAsu clone DmCD00765633 (Tempe, AZ, USA). The cDNA for TmodQ (NCBI Reference Sequence: NP_001263112.1) was synthesized by GeneWiz (South Plainfield, NJ, USA). Each cDNA was PCR-amplified and ligated between the SacI and KpnI sites of pRSF_Duet1 (Millipore Sigma, Burlington, MA, USA) with an N-terminal 6x histidine tag and a C-terminal Strep-II tag.

### Protein expression and purification

Tmod isoforms in pRSF_Duet1 were expressed in ArcticExpress(DE3) RIL cells (Agilent Technologies), grown in Terrific Broth (TB) medium for 6 h at 37 °C to an optical density of ∼1.5 at 600 nm (OD600), followed by 24 h at 10°C in with 0.4 mM isopropyl-β-D-thiogalactoside (IPTG). Cells were harvested by centrifugation (4,000g), resuspended in nickel buffer (20 mM HEPES pH 7.5, 300 mM NaCl, 10 mM imidazole, 1 mM PMSF), and lysed using a microfluidizer apparatus (Microfluidics). Proteins were first purified on nickel-NTA resin, washed with 20 column volumes of nickel buffer, and eluted using nickel buffer supplemented with 300mM imidazole. Elution fractions containing the Tmod proteins were then loaded onto Streptactin XT resin (IBA Life Sciences, Göttingen, Germany) and washed with 20 column volumes of strep buffer (20 mM HEPES pH 7.5, 150 mM NaCl, 1 mM EDTA). Proteins were eluted using strep buffer supplemented with 40mM biotin, and further purified through a SD200HL 26/600 gel filtration column (GE Healthcare) in strep buffer supplemented with 1 mM DTT. Human CapZ (Rao et al., 2014), rabbit skeletal muscle actin (Pardee et al., 1982), and rabbit skeletal muscle tropomyosin (Smillie, 1982) were purified as described.

### Pointed-end capping assay

Capping assays were carried out as described previously (Rao et al., 2014, Kumari et al., 2020). Briefly, 1.5 μM of pre-formed F-actin seeds, coated with 1 μM Tropomyosin and capped at the barbed end with saturating amounts of CapZ (25 nM), were incubated and assayed alone or with 4 different concentrations (25, 100, 400, 800 nM). Upon addition of 1.5 μM G-actin (6% pyrene-labeled), reactions were monitored as the fluorescence increase resulting from the incorporation of pyrene-actin monomers at the pointed-end. Pyrene fluorescence (excitation 365 nm, emission 407 nm) was detected using a plate reader (BioTek Cytation 5), and for each Tmod concentration four replicates were recorded. Maximum polymerization rates were determined by taking the first derivative of each curve, identifying the time interval corresponding to the greatest slope, and performing a linear regression on the original data within that time interval.

### Immunostaining and confocal imaging

Larvae were dissected in ice cold HL3.1 medium as described in (Brent et al., 2009) and fixed with 10% formalin (Sigma-Aldrich #HT501128) for 20 minutes. Larval filets were blocked with PBT-BSA [1x PBS supplemented with 0.3% Triton X-100 (Sigma-Aldrich #X100) and 0.1% BSA (Sigma-Aldrich #A7906)] for 30 minutes. Larval filets were then incubated with primary antibody (Table S5) overnight at 4°C, followed by washes in PBT-BSA. Incubation with Alexa Fluor-conjugated secondary antibodies (Table S5), Phalloidin and/or Hoechst 3342 were performed at room temperature for 2 hours, followed by washes in PBT. Subsequently, larval filets were mounted in ProLong Gold (Invitrogen #P36930) and slides were cured at room temperature for 24h. Z-stacks were acquired using a SP5 laser-scanning microscope (Leica Microsystems) with either dry objectives (HC PL APO CS 10x/0.40 and HC PL APO CS2 20x/0.75) or oil-immersion objectives (HC PL APO CS 20x/0.70, HCX PL APO CS 40x/1.25-0.75, HCX PL APO CS 63x/1.40-0.60, and HCX PL APO CS 100x/1.46). All samples intended for direct comparison were imaged using the same confocal settings. Maximum intensity projections of confocal Z stacks were rendered using Fiji. Approximately 7 larvae (range: 5-10) of each genotype were dissected and experiments were repeated at least twice.

### Quantification of muscle defects per segment

Muscle defects in second instar, early third instar, and late third instar larvae were scored based on ectopic and abnormal actin structures using a confocal microscope. A score was given for each VL3/4 muscle pair: 3 – WT-like, 2 – mild structural defects (diagonal myofibrils) and 1 – severe defects in muscle structure. We evaluated muscles in both hemisegments of the larva in abdominal segments 1-6. This analysis was performed in triplicate with 3-4 larvae per replicate.

### Sarcomere and thin filament size/count analysis

Larvae expressing a Zasp::GFP protein trap were fixed by brief (∼5 seconds) submersion in 65°C water and mounted on a slide with halocarbon oil. Z-stacks of the cuticular side of the VL3 or VL4 muscles from abdominal segment 2 were acquired using a SP5 laser-scanning microscope (Leica Microsystems) with either a HC PL APO CS2 20x/0.75 (late third instars) or HCX PL APO CS 40x/1.25-0.75 (second instars) oil immersion objective.

Muscle length, sarcomere size and number were quantified as described in step 32 of (Balakrishnan et al., 2021). For accuracy and to assess potential regional differences, sarcomere size was also quantified by an alternative method. Z-stacks were opened in Fiji and a line that spanned six Z-discs (five sarcomeres) was made using the line tool. Subsequently, the line intensity plot was obtained, and the X-coordinates (microns) of the peaks were acquired with the multi-point tool. The distance in microns between Z-disks was obtained by calculating the difference between the X-coordinates of the six peaks. This analysis was done in triplicate. In each replicate, six muscles were analyzed, each belonging to an individual larva. Three lines were traced per muscle: in the anterior fraction of the muscle, in the medial region and in the posterior end, with a total of ∼90 sarcomere size measurements per replicate (Fig. S4C).

Sarcomere length was obtained from Zasp::GFP heat-fixed larvae, which was performed by a direct method from Z-disc to Z-disc distance. Due to technical challenges in some areas of the muscles, two confirmatory methods were also employed. First, measurements were confirmed by comparing the heat-fixed samples with a formalin fixed sample. Second, we performed a sarcomere length average by taking the muscle length divided by the total number of Z-discs.

Thin filament length was assessed in formalin fixed samples that displayed a clear striation pattern and a clear M-line (therefore, we conclude the muscles were in a stretched configuration). Zasp was used as a marker for Z-discs, we traced a line spanning three contiguous Z-discs. The line intensity plot of Phalloidin was obtained and was normalized by calculating deviation from the mean to account for background noise; those with values above zero were used to calculate thin filament length.

### Intensity quantification at the muscle ends (MTJs)

Maximum intensity projections of confocal z-stacks of β_PS_-Integrin, Talin and pTyr immunostainings were generated in Fiji. A line spanning the MTJ and three contiguous sarcomeres (as a landmark reference) was drawn with the line tool. Line intensity plots were obtained with the intensity value for each pixel in the line. These values were split into five bins (each bin encompassing 20% of the intensity values) along the anterior-posterior axis and an average was calculated for each bin. Normalization was accomplished by calculating the deviation from the mean value. We traced two lines per MTJ per larva (16-20 measurements per genotype from 4-5 different larvae) and per genotype. This analysis was done in duplicate.

### Nuclear analysis

All images were processed and analyzed with Fiji. 2D quantifications of VL3 and VL4 muscles were performed with maximum intensity projections (as previously described in Windner et al. (2019)). Muscle areas were traced by hand with the polygon tool. Automated thresholding of fluorescence intensities of anti-Lamin and/or Hoechst was used to generate binary images of VL nuclei. The number, size (area), position (X and Y coordinates) and shape descriptors (roundness and circularity) of all nuclei within each cell was obtained and the binary masks were used to quantify Hoechst and H3K9ac pixel intensity (mean gray value). Automated thresholding of anti-Fibrillarin labeling was used to generate binary images of nucleoli and measure nucleolar sizes (areas) and to quantify Fibrillarin pixel intensity (mean gray value). The 3D rendering of nuclei was processed with Imaris.

### RNA sequencing analysis

Eight to ten late third instar larvae were dissected as described above per genotype in each replicate. The experiment was done in triplicate. The muscle-enriched larval filets were transferred to TRIzol and homogenized. The lysates were submitted to the MSKCC’s Integrated Genomics Operations (IGO) core for further total RNA extraction, quality assessment and Illumina next-generation sequencing. The sequencing libraries were constructed using SMARTerAmpSeq after the mRNAs were enriched by oligo-dT-mediated purification. The libraries were then sequenced on the Illumina HiSeq 4000 in a PE50 run. Over 40 million reads were generated per library. FASTQ data was read into R with the ShortRead *package (Morgan et al., 2009) and QC was performed with the Rfastp package (Wang et al., 2022). Reads were mapped and aligned to the Drosophila* genome using Rsubread (Liao et al., 2019). Next, the reads were counted with the GenomicAlignments package (Lawrence et al., 2013). Gene differential expression tests were performed using DESeq2 (Love et al., 2014). Finally, enrichment analysis was performed through goseq (Young et al., 2010) using both Gene Ontology and KEGG databases. Plots were drawn using R with the ggplot2 package (Wickham, 2016).

### ATP assay

An ATP calibration curve was performed using the stabilized ATP standard stocks provided in the kit ATP Bioluminescence Assay Kit HS II (Millipore Sigma #11699709001) and following the supplied protocol. Five to ten late third instar wandering larvae from each genotype were dissected (See Immunolabeling and Imaging) and transferred into the cell lysis reagent provided with the kit. Lysates were homogenized and protein concentrations were determined using the Bradford assay. Equal amounts of protein (10 μg) were used for bioluminescence measurements. Blank measurements were subtracted from the raw data and ATP concentrations were calculated from the standard curve data. The experiment was done in triplicate.

### Statistical analysis

Statistical significance was determined by either student’s unpaired t-test (comparisons between two groups), one-way ANOVA, or two-way ANOVA (comparisons among multiple groups) in the GraphPad Prism software. Data are presented as mean±SD, asterisks are used to denote significance. Graphs were made using either GraphPad Prism, Microsoft Excel or R.

## ACKNOWLEDGEMENTS

We thank the Baylies lab members, F. Schnorrer, G. Tanentzapf, A. Beggs and V. Gupta for helpful discussions and H. Bellen and F. Schöck for providing antibodies. We thank the members of the Integrated Genomics Operation (IGO) and the Bioinformatics Cores at MSKCC.

## DATA AVAILABILITY

RNA-seq data have been deposited in Gene Expression Omnibus (GEO) under the accession number GSE210507.

