## Supplemental information: 7 Supplemental Figures, 2 movie legends, 5 Supplementary Tables for "*Drosophila* Tropomodulin is required for multiple actin-dependent processes in myofiber assembly and maintenance"

### SUPPLEMENTARY FIGURES

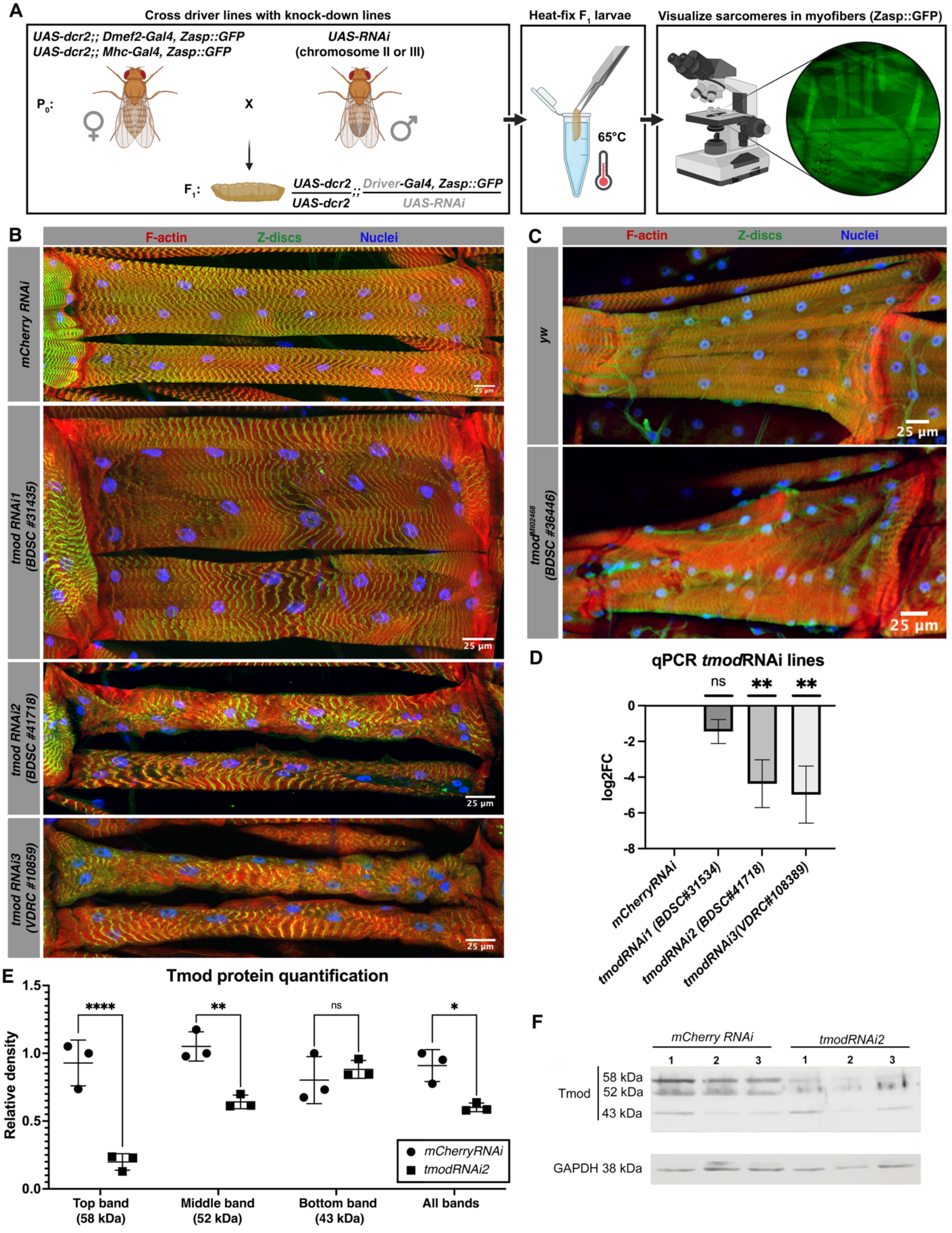


**Fig. S1: Confirmation of Tmod KD phenotype through multiple methods.**

(A) Muscle-specific KD screen set up showing the crosses between Driver-Gal4 lines and UAS-RNAi lines, heat fixation and visualization of the larval body-wall musculature structure through Zasp::GFP gene-trap.

(B) VL3 and VL4 myofibers (red, phalloidin; green, Zasp::GFP; blue, Hoechst) from segment A2 in control and three different Tmod RNAi lines displaying the same phenotype to different degrees at late third instar.

(C) VI1 muscle from segment A1 in *tmod*$^{MI02468}$ mutant and control myofibers at late third instar.

(D) Quantification of Tmod expression by qPCR in 3 different RNAi lines (N= 3 replicates, 7-10 larval carcasses per genotype per replicate; $p_{RNAi1}$=0.3119, $p_{RNAi2}$=0.0032, $p_{RNAi3}$=0.0014, ordinary one-way ANOVA multiple comparisons). Mean±SD.

(E and F) Western blot and quantification of Tmod in late third instar muscle-enriched lysates. Glyceraldehyde 3-phosphate dehydrogenase (GAPDH) is used as a loading control. Normalized to GAPDH. Each dot is a replicate (N=3 replicates, n=5-10 larval carcasses per genotype per experiment, $p_{Top\ band}$<0.0001, $p_{Middle\ band}$=0.0012, $p_{Bottom\ band}$=0.8602 , $p_{All\ bands}$=0.0130, ordinary 2-way ANOVA multiple comparisons). Mean±SD.

Scale bars: 25μm (A and B)

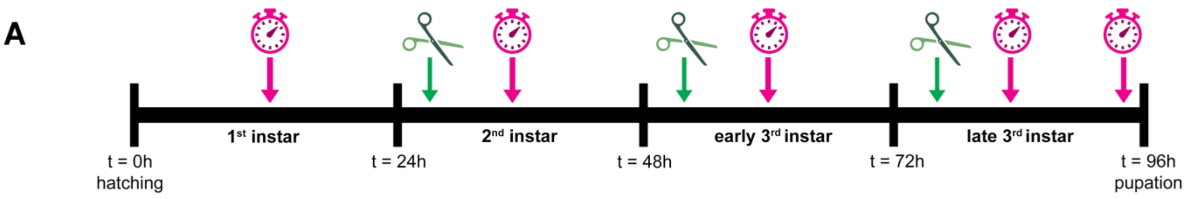


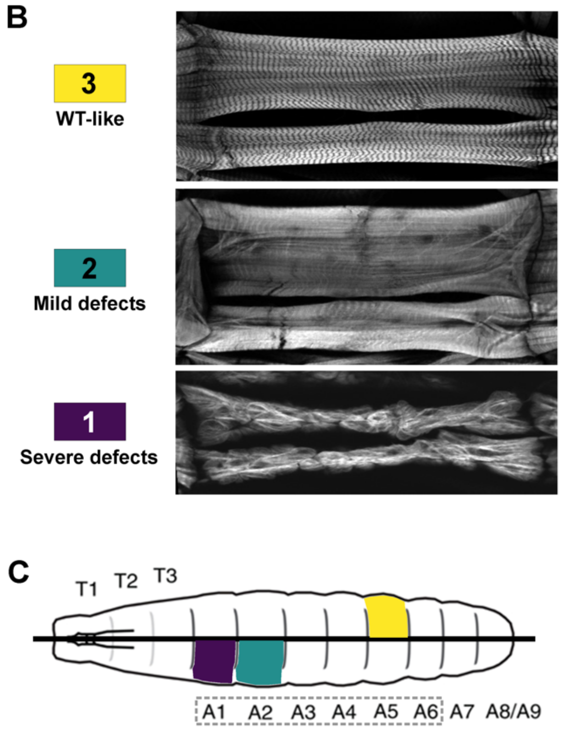


**D** Quantification of affected segments by stage

| | A1 | A2 | A3 | A4 | A5 | A6 | Avg. score |
|---|---|---|---|---|---|---|---|
| 2nd instar | 2 | 2.5 | 3 | 3 | 3 | 3 | 2.8 |
| Early 3rd instar | 1.3 | 1.9 | 2.2 | 2.7 | 3 | 3 | 2.4 |
| Late 3rd instar | 1 | 1.2 | 1.9 | 2.2 | 2.5 | 2.6 | 1.9 |

**E** Quantification of affected segments by RNAi line

| | A1 | A2 | A3 | A4 | A5 | A6 | Avg. score |
|---|---|---|---|---|---|---|---|
| mCherry | 3 | 3 | 3 | 3 | 3 | 3 | 3 |
| tmod RNAi1 | 2 | 2 | 3 | 3 | 3 | 3 | 2.7 |
| tmod RNAi2 | 1 | 1 | 1.4 | 2 | 2.3 | 2.125 | 1.6 |
| tmod RNAi3 | 1 | 1 | 1 | 1 | 1 | 1 | 1 |

**Fig. S2: Experimental design and quantification of myofibril defects.**

(A) Experimental setup indicating when larval dissection (scissor symbol, green) or locomotion assays (timer symbol, magenta) were performed during the larval stages.

(B) Scoring system (3, yellow, WT-like; 2, aquamarine, mild defects and 1, purple, severe defects) for F-actin (gray, phalloidin) defects in assessed (VL3 and VL4) muscles.

(C) Larval diagram showing hemisegments T1-A9. Colored polygons display example scoring. Gray dotted line indicates quantified hemisegments.

(D and E) Quantification of myofibril (F-actin) defects in V1 or VL3/VL4 in segments A1-A6 by stage (from second instar to late third instar) or by RNAi line in late third instar (N = 3 replicates, n= 5 larvae per genotype per stage).

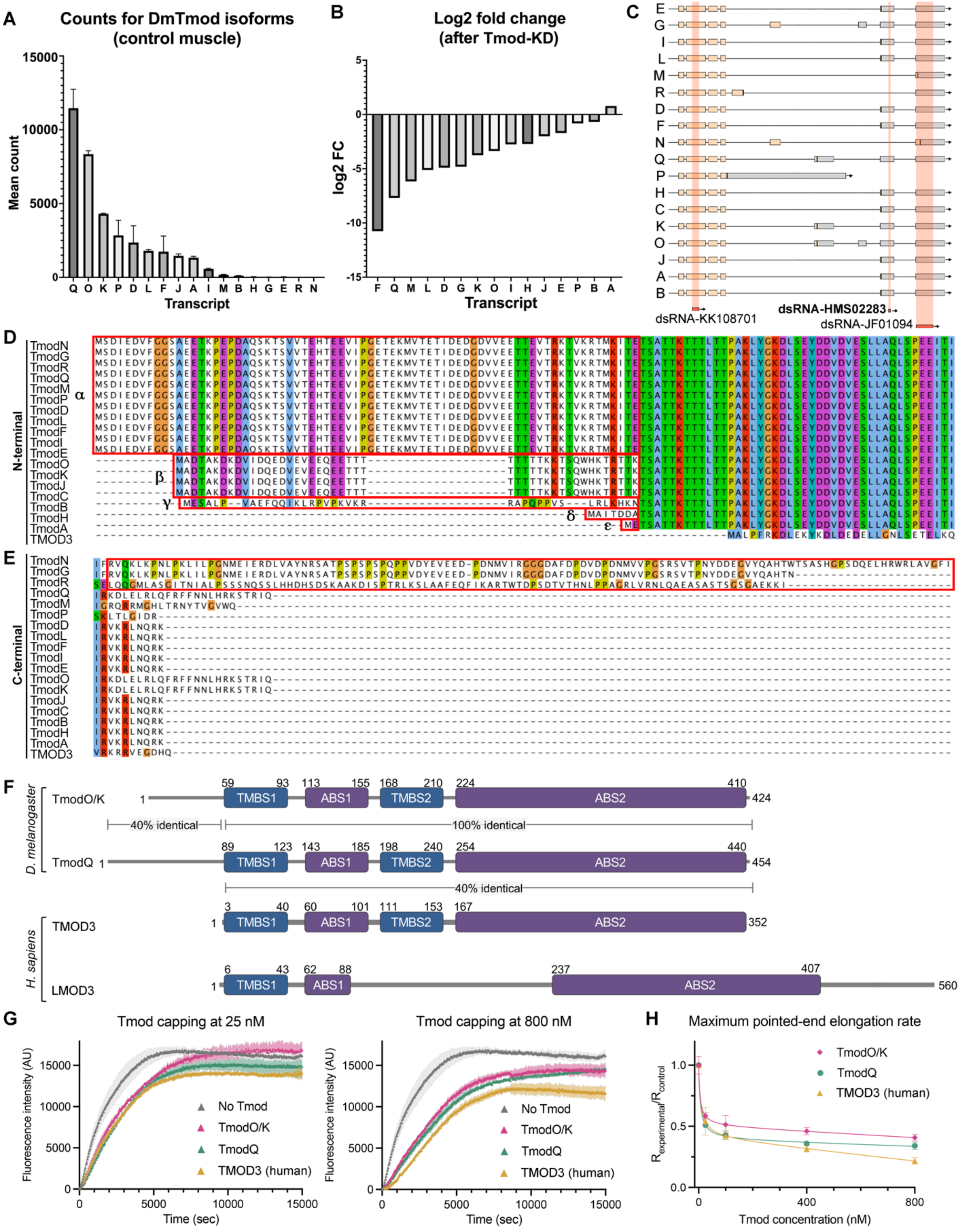


**Fig. S3: Analysis of *Drosophila* Tmod expression, structure, and function.**

(A) Normalized counts average from bulk, muscle-enriched RNAseq data for the different *Drosophila* Tmod transcripts in control larvae. Isoforms C, E, G, H, N, R were either non-detected or expressed at low levels in control larvae. Mean±SD.

(B) Log2 fold change of Tmod transcript expression in *tmod* KD vs control from muscle-enriched RNAseq data ($p_F$<0.0001, $p_Q$<0.0001, $p_M$<0.0001, $p_L$<0.0001, $p_D$<0.0001, $p_G$=0.0002, $p_K$<0.0001, $p_O$<0.0001, $p_I$<0.0001, $p_H$=0.2868, $p_J$<0.0001, $p_E$=0.8967, $p_P$=0.0028, $p_B$=0.7599, $p_A$=0.0024). The Tmod-KD construct used for the main experiments (dsRNA-HMS02283) reduced levels of most Drosophila Tmod isoforms except isoforms A, B and P. RNAseq data was used for plots A, B (N=3 replicates with n=7-10 larval carcasses per genotype per N).

(C) Schematic of 18 different C-terminal Tmod transcripts and RNAi targeting region. The Tmod-KD construct used for the main experiments (dsRNA-HMS02283) targeted all isoforms (either at the 3'-UTR region or at the last intronic region) except isoform P. Beige boxes indicate coding regions, grey boxes indicate untranslated regions, dark orange boxes and transparent red panels indicate where the dsRNA construct targets.

(D) Amino acid sequence alignment of *Drosophila* Tmod isoforms and human TMOD3 at the N-terminal region. Based on their N-termini extensions, the 18 isoforms can be classified into four subgroups: α displays extensions of 88 amino acids (D, E, F, G, I, L, M, N, P, Q, R), β displays extensions of 58 amino acids (C, J, K, O), γ displays extensions of 52 amino acids (B), and δ/ε (A, H) which lack an N-terminal extension.

(E) Amino acid sequence alignment of *Drosophila* Tmod isoforms and human TMOD3 at the C-terminal region. Three of the isoforms (red box) have Proline-rich C-terminal extensions (G, N, R), but these are expressed at low levels and were not further considered. Thus, we focused on isoforms differing at the N-terminus.

(F) Domain maps of and similarity of *Drosophila* and Human TMOD/LMOD genes. TMBS = tropomyosin binding site, ABS = actin binding site. Sequence analysis further showed that all the *Drosophila* Tmod isoforms share an identical central region, displaying strong similarity with mammalian Tmods. This region comprises all domains implicated in pointed-end capping, including two Tropomyosin- and two actin-binding sites (Rao et al., 2014).

(G) Pyrene-labeled actin polymerization assays comparing capping efficiencies at the pointed end with no Tmod, human TMOD3, *Drosophila* TmodO/K and *Drosophila* TmodQ at 25nM and 800nM. Data is from Fig. 3A (N=4 replicates, per isoform and per concentration). Mean±SD (lighter-color shading).

(H) Maximum pointed end elongation rate (R) from the data in Fig. 3A displaying greatest slope in the curves (TMOD3, Tmod O/K and TmodQ) normalized to the slope without Tmod. Lower R values indicate slower elongation i.e., stronger capping activity. Mean±SD.

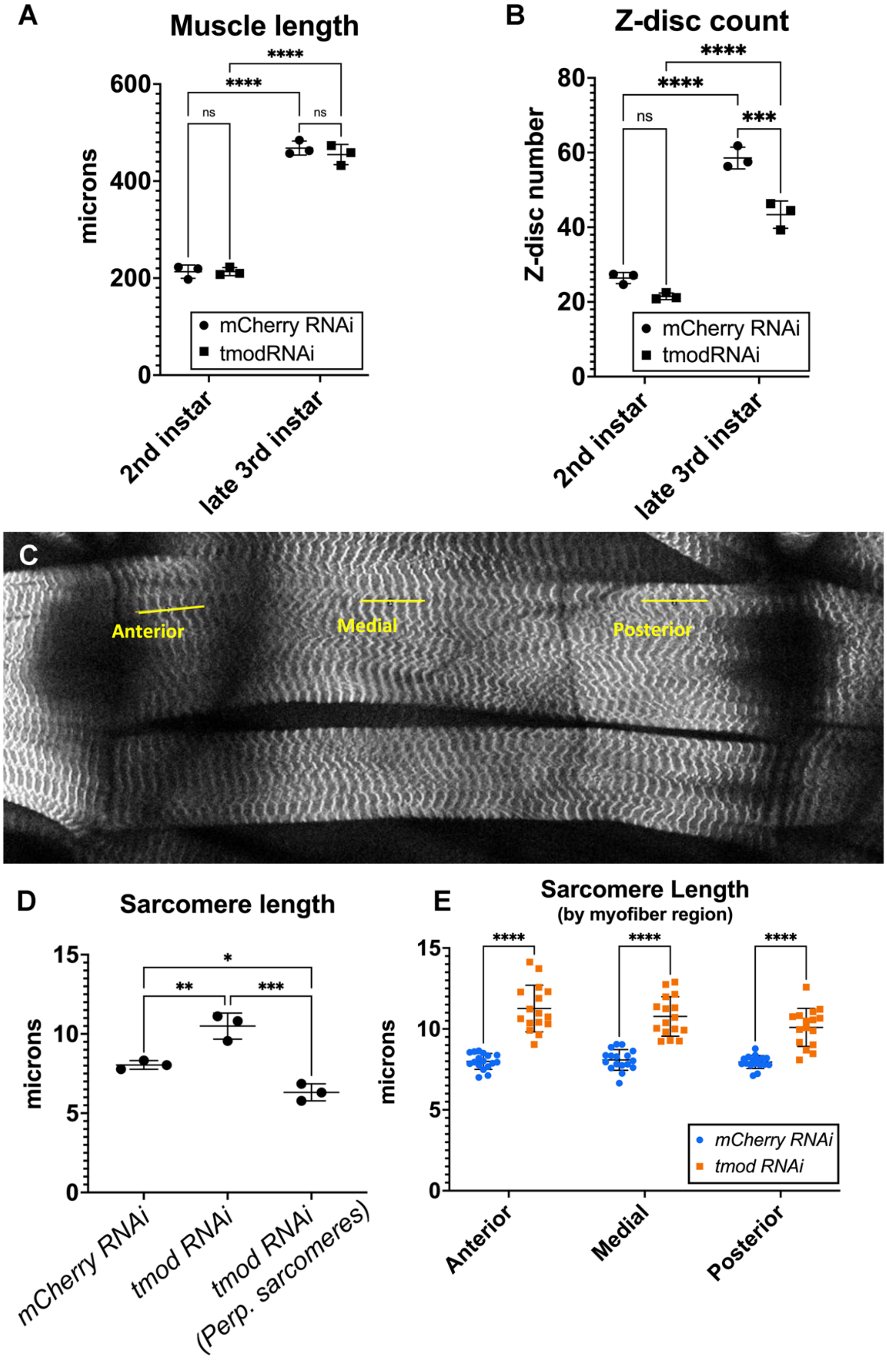


**Fig. S4: Analysis of multiple sarcomere-related parameters in VL3 myofibers from segment A2.**

(A) Quantification of average muscle length in second and late third instars in control and *tmod* KD. Each dot is one replicate N=3 replicates with $n_{2nd\ instars}$=19-20, $n_{late\ 3rd\ instars}$=16-17 myofibers per genotype; $p_{2nd\ instars}$>0.9999, $p_{3rd\ instars}$=0.8893, $p_{mCherryRNAi}$<0.0001, $p_{tmodRNAi}$<0.0001, ordinary 2-way ANOVA multiple comparisons.

(B) Quantification of average Z-disc count in second and late third instars in control and *tmod* KD. Each dot is one replicate, N=3 replicates with $n_{2nd\ instars}$=19-20, $n_{late\ 3rd\ instars}$=16-17 myofibers per genotype; $p_{2nd\ instars}$=0.1537, $p_{3rd\ instars}$=0.0003, $p_{mCherryRNAi}$<0.0001, $p_{tmodRNAi}$<0.0001, ordinary 2-way ANOVA multiple comparisons.

(C) VL3 and VL4 muscles (white, Zasp::GFP) in segment A2 of a control larva. Yellow lines show the regionalization of the six Z-disc sets used for the quantification in E.

(D) Average sarcomere length quantification in late third instar comparing control, *tmod* KD and perpendicular *tmod* KD sarcomeres. Each dot is one replicate: N=3 replicates with n=45-105 sarcomeres per genotype per dot [from 3-7 different larvae], $n_{perp}$=9-45 sarcomeres per dot; $p_{tmodRNAi\ vs\ mCherry\ RNAi}$=0.0054, $p_{Perp\ vs\ mCherryRNAi}$=0.0268, $p_{tmodRNAi\ vs\ perp.}$=0.0003, ordinary one-way ANOVA multiple comparisons.

(E) Average sarcomere length quantification by region in the muscle as shown in C. Each dot is one larva [16-17 larvae] (N = 3 replicates, n=5 sarcomeres per dot, per genotype; $p_{anterior}$<0.0001 , $p_{middle}$<0.0001, $p_{posterior}$<0.0001, ordinary 2-way ANOVA multiple comparisons). Mean±SD.

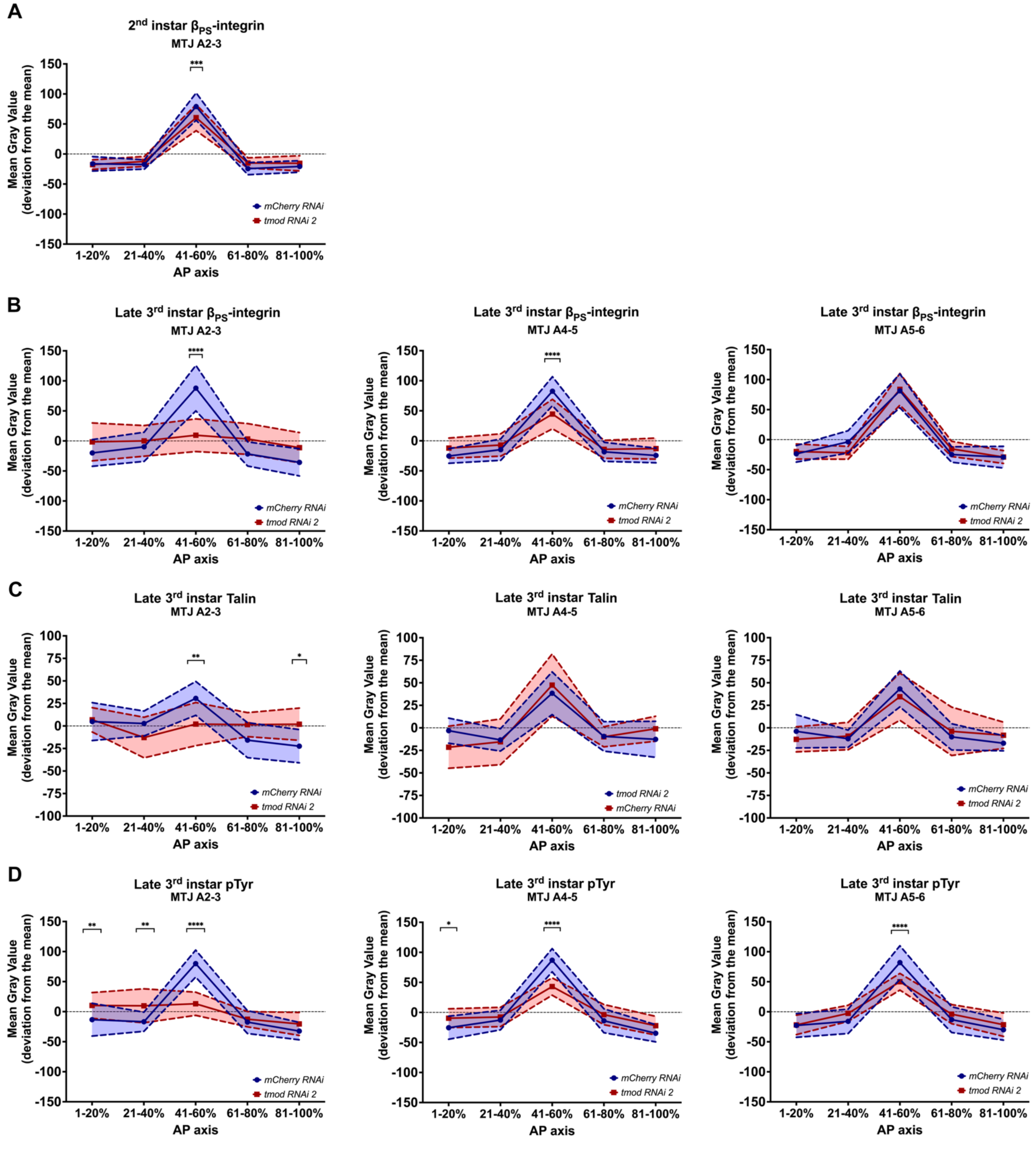


**Fig. S5: Quantification of tension-mediating proteins and pTyr at the MTJ.**

(A) Normalized $\beta_{PS}$-integrin pixel intensity signal along the MTJ (segments A2-3) in control (blue) and *tmod* RNAi (red) in second instar larvae. Graph shows one experiment (N>2 replicates, n=16-20 measurements per genotype [from 4-5 different larvae]; $p_{41\text{-}60\%}$=0.0003, ordinary 2-way ANOVA multiple comparisons).

(B-D) Normalized $\beta_{PS}$-integrin, Talin and pTyr pixel intensity signal along the MTJ (left, segments A2-3; middle, segments A4-5; right, segments A5-6) in control and *tmod* RNAi in wandering late third instar larvae. Graph shows one experiment (N > 2 replicates, n=12 measurements per genotype and per segment [from 6 different larvae]; ordinary 2-way ANOVA multiple comparisons). Transparent zones show SD for all graphs.

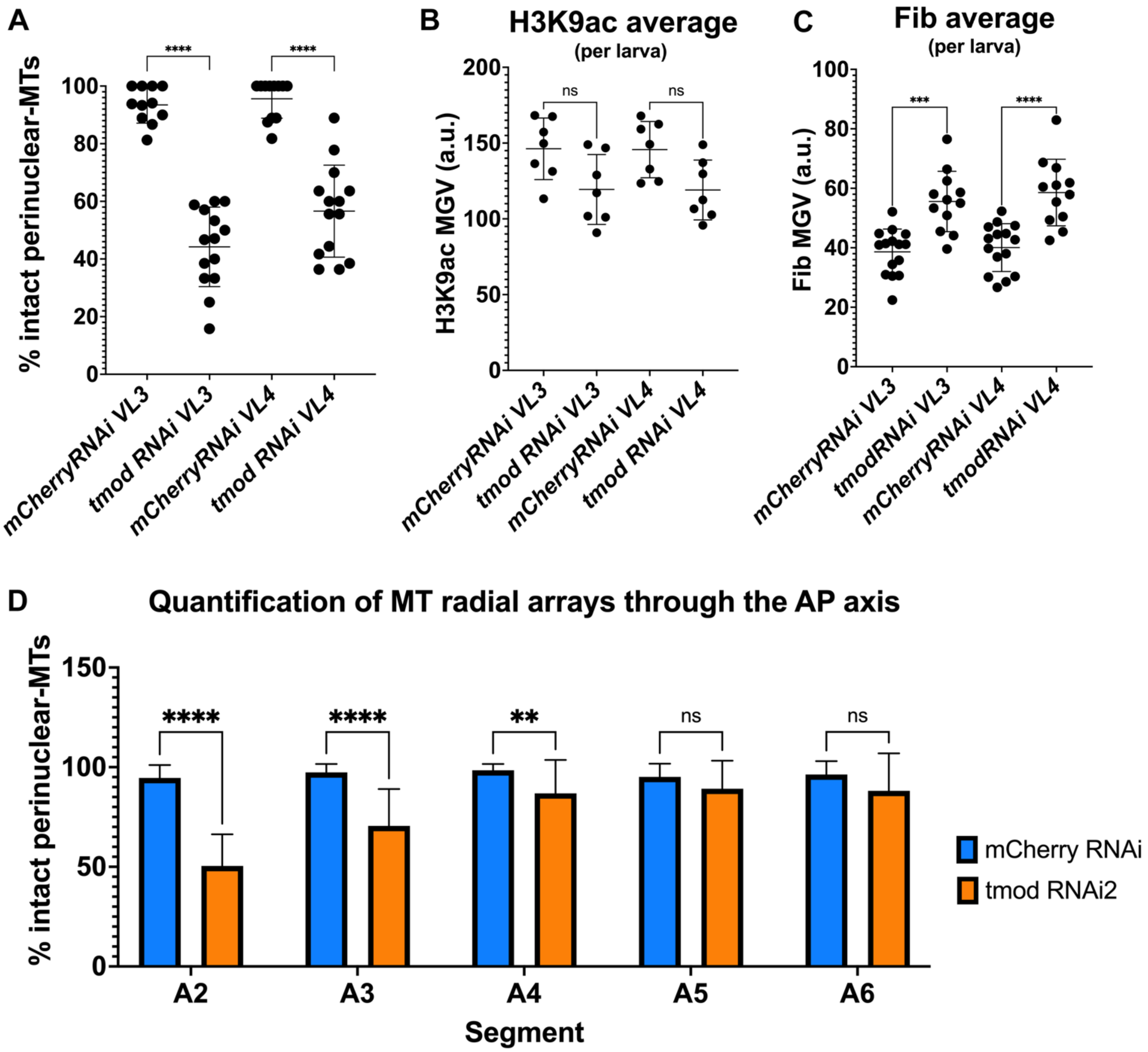


**Fig. S6: *tmod* KD VL3 and VL4 myofibers of segment A2 display defects in nuclear supporting structures and nuclear output averages in late third instars.**

(A) Quantification of intact perinuclear-MT radial arrays in control and *tmod* KD myofibers. Graph represents one experiment, each dot is one myofiber ($n_{VL3}$=11-14, $n_{VL4}$=11-14 myofibers [N=6 larvae]; $p_{VL3}$<0.0001, $p_{VL4}$<0.0001; ordinary one-way ANOVA multiple comparisons).

(B) Average pixel intensity quantification of nuclear H3K9ac in control and *tmod* KD myofibers. Graph represents one experiment, each dot is one myofiber per larva ($n_{VL3}$=7, $n_{VL4}$=7 myofibers; $p_{VL3}$=0.0936, $p_{VL4}$=0.0971; ordinary one-way ANOVA multiple comparisons).

(C) Average pixel intensity quantification of nuclear Fib in control and *tmod* KD myofibers. Graph represents one experiment, each dot is one myofiber ($n_{VL3}$=12-15, $n_{VL4}$=12-15 myofibers [from 7-8 larvae]; $p_{VL3}$=0.0001, $p_{VL4}$<0.0001; ordinary one-way ANOVA multiple comparisons).

(D) Quantification of intact perinuclear-MT radial arrays in control and *tmod* KD myofibers through the AP axis. Graph represents one experiment ($n_{A2}$=23-28, $n_{A3}$=23-30, $n_{A4}$=24-28, $n_{A5}$=22-32, $n_{A6}$=22-31 myofibers [mixed VL3 and VL4; from 6 larvae]; $p_{A2}$<0.0001, $p_{A3}$<0.0001, $p_{A4}$=0.0096, $p_{A5}$=0.5246, $p_{A6}$=0.1378, ordinary two-way ANOVA multiple comparisons).

Experiments repeated N>2. Mean±SD.

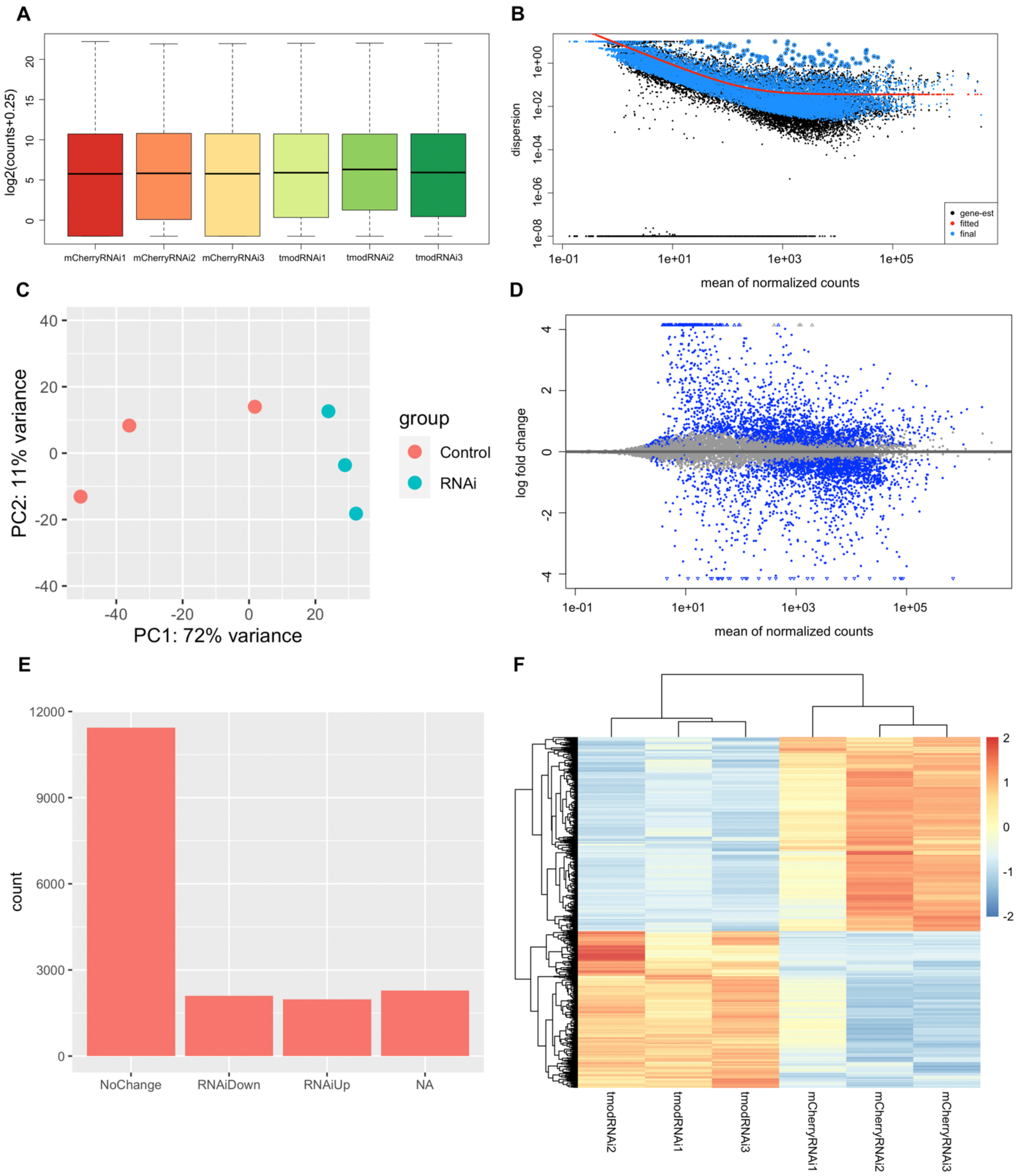


**Fig. S7: Analysis and verification of count data from RNAseq in control and *tmod* KD muscle-enriched late third instar carcasses.** (A) Boxplots depicting DESeq2 normalization of RNAseq samples [log2(counts+0.25)]. (B) Dispersion estimates plot showing variance/mean relationship and shrinkage (blue dots, adjusted dispersions; black dots, original dispersions; red line, dispersion-mean trend; black dots circled in blue, dispersion outliers not shrunk). (C) Principal component analysis (PCA) plot showing samples in 2D plane (two principal components) using all genes. (D) MA-plot displaying relationship between shrunken log2 fold-changes and expression levels (blue, genes with a $p_{adj}<0.05$). (E) Plot displaying number of genes (count) that do not show a change in expression ($p_{adj}>0.05$), number of genes downregulated ($p_{adj}<0.05$, log2FC<0) and number of genes upregulated ($p_{adj}<0.05$, log2FC>0). (F) Heatmap of relative changes in gene expression using Z-score for each row (gene). Only 2526 genes are shown ($p_{adj}<0.01$) in Tmod KD muscle. The color scale indicates the gene expression standard deviations from the mean within each row (blue, low relative expression; red, high relative expression).

**SUPPLEMENTARY MOVIES**

Movie 1: 3D rendering of disc-shaped control myonuclei at high magnification at the visceral side of the myofiber (green, Phalloidin; red, Lamin).

Movie 2: 3D rendering of misshapen and internalized myonuclei at high magnificationin the Tmod KD myofiber (green, Phalloidin; red, Lamin).

### SUPPLEMENTARY TABLES

#### Table S1: Summary of tested lines.

| Fly potential orthologues | Human gene | No. RNAi variants |
|---|---|---|
| Kelch (CG7210) | KBTBD13, KLHL40, KLHL41 | 3 |
| Lasp (CG3849) | Nebulin | 4 |
| Mhcl (CG31045) | Myosin XVIII B | 2 |
| Slim (CG5186) | KBTBD13, KLHL40, KLHL41 | 4 |
| Tm1 (CG4898) | Tropomyosin 2/3 | 1 |
| Tm2 (CG4843) | Tropomyosin 2/3 | 5 |
| Tmod (CG1539) | Leiomodin 3 | 4 |
| Upheld (CG7107) | Troponin T1 | 3 |
| CG-lines (with Kelch and BTB domains) | KBTBD13, KLHL40, KLHL41 | 3 |
| | **Total No. RNAi tested** | 29 |

#### Table S2: Phenotypes per genotype in larvae, pupae, and adults.

| ***Drosophila* gene** | **Catalog number and genotype** | **UAS-dcr2;; Mhc-Gal4, Zasp::GFP** | | | **UAS-dcr2;; Dmef2-Gal4, Zasp::GFP** | | |
|---|---|---|---|---|---|---|---|
| | | **Larvae** | **Pupae** | **Adults** | **Larvae** | **Pupae** | **Adults** |
| **Kelch (CG7210)** | BDSC#31251 y[1] v[1]; P{y[+t7.7] v[+t1.8]=TRiP.JF01768}attP2 | ✓ | ✓ | ✓ | ✓ | ✓ | ✗ (parachute) |
| | BDSC#55612 y[1] sc[*] v[1]; P{y[+t7.7] v[+t1.8]=TRiP.HMC03751}attP40 | ✓ | ✓ | ✓ | ✓ | ✓ | ✓ |
| | VDRC#105397 P{KK100891}VIE-260B | ✓ | ✓ | ✓ | ✓ | ✗ (die late pupae) | - |
| **Lasp (CG3849)** | BDSC#26305 y[1] v[1]; P{y[+t7.7] v[+t1.8]=TRiP.JF02075}attP2 | ✓ | ✓ | ✓ | ✓ | ✓ | ✗ (parachute) |
| | VDRC#47126 w[1118] P{GD16308}v47126 | ✓ | ✓ | ✓ | ✓ | ✗ (die early pupae) | - |

| | | | | | | | |
|---|---|---|---|---|---|---|---|
| | VDRC#47127<br>w[1118] P{GD16308}v47127 | ✓ | ✓ | ✓ | ✓ | ✗<br>(die early pupae) | - |
| | VDRC#109416<br>P{KK108117}VIE-260B | ✓ | ✓ | ✓ | ✓ | ✗<br>(die early pupae) | - |
| **Mhcl (CG31045)** | BDSC#51456<br>y[1] v[1]; P{y[+t7.7]<br>v[+t1.8]=TRiP.HMC03191}attP40 | ✓ | ✓ | ✓ | ✓ | ✓ | ✓ |
| | BDSC#62973<br>y[1] v[1]; P{y[+t7.7]<br>v[+t1.8]=TRiP.HMJ30050}attP40 | ✓ | ✓ | ✓ | ✓ | ✓ | ✓ |
| **Slim (CG5186)** | BDSC#29437<br>y[1] v[1]; P{y[+t7.7]<br>v[+t1.8]=TRiP.JF03373}attP2 | ✓ | ✓ | ✓ | ✓ | ✓ | ✓ |
| | BDSC#34376<br>y[1] sc[*] v[1]; P{y[+t7.7]<br>v[+t1.8]=TRiP.HMS01366}attP2 | ✓ | ✓ | ✓ | ✓ | ✓ | ✓ |
| | VDRC#15185<br>w[1118]; P{GD4969}v15185 | ✓ | ✓ | ✗<br>(flightless) | ✗<br>(embryonic lethal) | - | - |
| | VDRC#108067<br>P{KK107448}VIE-260B | ✓ | ✓ | ✓ | ✓ | ✗<br>(die late pupae) | - |
| **Tm1 (CG4898)** | VDRC#34119<br>w[1118]; P{GD10529}v34119 | ✓ | ✓ | ✓ | ✓ | ✗<br>(die early pupae) | - |
| **Tm2 (CG4843)** | BDSC#31535<br>y[1] v[1]; P{y[+t7.7]<br>v[+t1.8]=TRiP.JF01095}attP2 | ✓ | ✓ | ✗<br>(flightless) | ✗<br>(embryonic lethal) | - | - |
| | BDSC#41695<br>y[1] sc[*] v[1]; P{y[+t7.7]<br>v[+t1.8]=TRiP.HMS02260}attP2 | ✓ | ✓ | ✗<br>(parachute<br>) | ✓ | ✓ | ✗<br>(flightless) |
| | VDRC#42008<br>w[1118]; P{GD11445}v42008 | ✓ | ✓ | ✗<br>(flightless) | ✗<br>(embryonic lethal) | - | - |
| | VDRC#42010<br>w[1118]; P{GD11445}v42010 | ✓ | ✓ | ✗<br>(flightless) | ✗<br>(embryonic lethal) | - | - |
| | VDRC#107970<br>P{KK111307}VIE-260B | ✓ | ✓ | ✗<br>(flightless) | ✗<br>(embryonic lethal) | - | - |
| **Tmod (CG1539)** | BDSC#31534<br>y[1] v[1]; P{y[+t7.7]<br>v[+t1.8]=TRiP.JF01094}attP2 | ✓ | ✓ | ✓ | ✓ | ✓ | ✗<br>(parachute) |
| | BDSC#41718<br>y[1] sc[*] v[1]; P{y[+t7.7]<br>v[+t1.8]=TRiP.HMS02283}attP2 | ✓ | ✓ | ✓ | ✗<br>(thin muscles) | ✗<br>(few late pupae die) | ✓<br>(the ones that survive) |
| | VDRC#32602<br>w[1118]; P{GD9005}v32602 | ✓ | ✓ | ✓ | N/A | N/A | N/A |
| | VDRC#108389<br>P{KK108701}VIE-260B | ✓ | ✓ | ✓ | ✗<br>(thin muscles) | ✗<br>(die early pupae) | - |
| **Upheld (CG7107)** | BDSC#31541<br>y[1] v[1]; P{y[+t7.7]<br>v[+t1.8]=TRiP.JF01102}attP2 | ✓ | ✓ | ✗<br>(flightless) | ✗<br>(embryonic lethal) | - | - |
| | BDSC#32949<br>y[1] sc[*] v[1]; P{y[+t7.7]<br>v[+t1.8]=TRiP.HMS00743}attP2 | ✗<br>(forever $2^{nd}$ instar) | - | - | ✗<br>(embryonic lethal) | - | - |
| | VDRC#27853<br>w[1118]; P{GD12132}v27853 | ✓ | ✓ | ✗<br>(flightless) | ✗<br>(embryonic lethal) | - | - |
| **CG1812** | BDSC#38974<br>y[1] v[1]; P{y[+t7.7]<br>v[+t1.8]=TRiP.HMS01890}attP40 | ✓ | ✓ | ✓ | ✓ | ✓ | ✓ |

| | | | | | | | |
|---|---|---|---|---|---|---|---|
| | BDSC#42793<br>y[1] v[1]; P{y[+t7.7]<br>v[+t1.8]=TRiP.GL01163}attP2 | ✓ | ✓ | ✓ | ✓ | ✗<br>(die early pupae) | ✓ |
| **CG1126** | BDSC#65367<br>y[1] sc[*] v[1]; P{y[+t7.7]<br>v[+t1.8]=TRiP.HMC06119}attP40 | ✓ | ✓ | ✓ | ✓ | ✓ | ✓ |

**Table S3:** ***tmod*** **transcript expression (mean scaled counts) and log2 fold change (log2FC) in control and Tmod KD muscles from RNAseq.**

| Transcript (Flybase identifier) | Control larvae (mean scaled counts) | Tmod KD larvae (mean scaled counts) | Log2FC | Padj |
|---|---|---|---|---|
| A (FBtr0085662) | 1335.49 | 2295.18 | 0.78 | 2.43E-03 |
| B (FBtr0085663) | 111.00 | 68.93 | -0.69 | 7.60E-01 |
| D (FBtr0085659) | 2359.63 | 78.33 | -4.91 | 8.91E-52 |
| E (FBtr0085658) | 22.31 | 6.83 | -1.71 | 8.97E-01 |
| F (FBtr0085660) | 1736.46 | 1 | -10.76 | 1.73E-85 |
| G (FBtr0113483) | 28.03 | 1 | -4.81 | 1.97E-04 |
| H (FBtr0302110) | 42.63 | 6.44 | -2.73 | 2.87E-01 |
| I (FBtr0302111) | 569.15 | 83.83 | -2.76 | 4.22E-11 |
| J (FBtr0302112) | 1459.62 | 361.86 | -2.01 | 3.17E-14 |
| K (FBtr0302600) | 4307.70 | 317.28 | -3.76 | 3.15E-21 |
| L (FBtr0302601) | 1804.21 | 51.82 | -5.12 | 5.99E-40 |
| M (FBtr0306685) | 174.20 | 2.42 | -6.17 | 3.36E-08 |
| N (FBtr0306686) | 9.54 | 0.68 | -3.80 | N/A (below count threshold) |
| O (FBtr0306687) | 8348.16 | 808.00 | -3.37 | 4.44E-72 |
| P (FBtr0336446) | 2840.16 | 1600.08 | -0.83 | 2.83E-03 |

| | | | | |
|---|---|---|---|---|
| Q (FBtr0336447) | 11460.57 | 55.64 | -7.69 | 1.14E-34 |
| R (FBtr0479778) | 11.93 | 3.15 | -1.92 | N/A (below count threshold) |

**Table S4: Actin isoform expression (mean scaled counts) and log2 fold change (log2FC) in control and Tmod KD muscles from RNAseq.**

| Gene symbol (Flybase identifier) | Control larvae (mean scaled counts) | Tmod KD larvae (mean scaled counts) | Log2FC | Padj |
|---|---|---|---|---|
| Act57B (FBgn0000044) | 1054400.49 | 3593787.25 | 1.77 | 5.30E-03 |
| Act87E (FBgn0000046) | 80487.48 | 378031.68 | 2.23 | 3.37E-07 |
| Act5C (FBgn0000042) | 306248.53 | 178954.30 | -0.78 | 1.14E-03 |
| Act42A (FBgn0000043) | 55923.02 | 29806.44 | -0.91 | 4.88E-06 |

**Table S5: Key resources table.**

| REAGENT or RESOURCE | SOURCE | IDENTIFIER |
|---|---|---|
| **Antibodies** | | |
| Chicken anti-GFP (IF 1:200) | Abcam | Cat#13970; RRID: AB_300798 |
| Rabbit anti-Zasp (IF 1:400) | From F. Schöck (McGill) | Jani K. and Schöck F., 2007 |
| Mouse anti-$\beta_{PS}$-Integrin (Mys) (IF 1:50) | Developmental Studies Hybridoma Bank | Cat#CF.6G11; RRID: AB_528310 |
| Mouse anti-Talin (Rhea) (IF 1:1 mix of each clone, final concentration of 1:5) (WB :1 mix of each clone, final concentration of 1:100; 5% milk) | Developmental Studies Hybridoma Bank | Cat#A22A; RRID: AB_10660289<br>Cat#E16B; RRID: AB_10683995 |
| Mouse anti-pTyr (clone 4G10) (IF 1:500) | Millipore Sigma | Cat#05-321; RRID: AB_309678 |
| Mouse anti-Lamin (IF 1:50) | Developmental Studies Hybridoma Bank | Cat#ADL67.10; RRID: AB_528336 |
| Mouse anti-Fibrillarin (IF 1:100) | EnCor Biotechnology | Cat#MCA-38F3; RRID: AB_2278545 |

| | | |
|---|---|---|
| Rabbit anti-H3K9ac (IF 1:200) | Active Motif | Cat#39137; RRID: AB_2561017 |
| Mouse anti-alpha tubulin (clone DM1A) (IF 1:500) | Millipore Sigma | Cat#T6199; RRID: AB_212802 |
| Guinea pig anti-Tmod (IF and WB 1:1,000) | From H. Bellen (Baylor College of Medicine) | Lighthouse D. et al., 2008 |
| Mouse anti-GAPDH (WB 1:5,000; 5% BSA) | Abcam | Cat#ab9484; RRID: AB_307274 |
| Alexa Fluor 405, 488, 555, 647 Phalloidin (IF 1:200) | Thermo Fisher | Cat#A30104<br>Cat#A12379<br>Cat#A34055<br>Cat#A22287; RRID: AB_2620155 |
| Hoechst 33342 | Thermo Fisher | Cat#H3570; RRID: AB_10626776 |
| Alexa Fluor 488, 555, 647 conjugated fluorescent secondary antibodies (1:200) | Thermo Fisher | N/A |
| Peroxidase-conjugated donkey anti-mouse (WB 1:5,000) | Jackson Immunoresearch | Cat#715-035-151; RRID: AB_2340771 |
| Peroxidase-conjugated donkey anti-guinea pig (1:5,000) | Jackson Immunoresearch | Cat#706-035-148; RRID: AB_2340447 |
| **Experimental Models: Organisms/Strains** | | |
| *D. melanogaster: yw* | Bloomington Drosophila Stock Center | BDSC#1495 |
| *D. melanogaster: UAS-dicer2;; DMef2-Gal4, Zasp66::GFP* | This study | N/A |
| *D. melanogaster: UAS-dicer2;; Mhc-Gal4, Zasp66::GFP* | This study | N/A |
| *D. melanogaster: UAS-mCherry-RNAi* | Bloomington Drosophila Stock Center | BDSC#35785 |
| *D. melanogaster: UAS-tmod-RNAi2* (dsRNA-HMS02283) | Bloomington Drosophila Stock Center | BDSC#41718 |
| *D. melanogaster: UAS-tmod-RNAi1* (dsRNA-JF01094) | Bloomington Drosophila Stock Center | BDSC#31534 |
| *D. melanogaster: UAS-tmod-RNAi3* (dsRNA-KK108701) | Vienna Drosophila Resource Center | VDRC#108389 |
| *D. melanogaster: tmod$^{MI0246}$* | Bloomington Drosophila Stock Center | BDSC#36446 |

| | | |
|---|---|---|
| **Oligonucleotides** | | |
| Primer: Forward Primer (5' -> 3') to Drosophila tmod: “TCCCGATGACAACTTCCTGC” | DRSC/TRiP Functional Genomic Resources | N/A |
| Primer: Reverse Primer (5' -> 3') to Drosophila tmod: “ CGATGGCCTGCTTATTGATGT” | DRSC/TRiP Functional Genomic Resources | N/A |
| Primer: Forward Primer (5' -> 3') to Drosophila Rpl32: “TGGTTTCCGGCAAGCTTCAA” | This study | N/A |
| Primer: Reverse Primer (5' -> 3') to Drosophila Rpl32: “ TGTTGTCGATACCCTTGGGC ” | This study | N/A |
| **Software** | | |
| ImageJ/Fiji | NIH Image | https://fiji.sc; RRID:SCR_002285 |
| GraphPad Prism | GraphPad | https://www.graphpad.com; RRID:SCR_002798 |
| Excel | Microsoft | https://products.office.com/en-us/excel; RRID:SCR_016137 |
| Imaris | Bitplane | http://www.bitplane.com/imaris/imaris; RRID:SCR_007370 |
| R | R Foundation for Statistical Computing | https://www.r-project.org; RRID:SCR_001905 |
| Bioconductor | Bioconductor | http://www.bioconductor.org; RRID:SCR_006442 |
| Photoshop | Adobe | https://www.adobe.com/products/photoshop.html; RRID:SCR_014199 |
| Illustrator | Adobe | https://www.adobe.com/products/illustrator.html; RRID:SCR_010279 |
| BioRender | Biorender | http://biorender.com; RRID:SCR_018361 |
